# Assessing nonlinear associations between body mass index and brain microstructure across adolescence

**DOI:** 10.64898/2026.04.02.716201

**Authors:** Alison Rigby, Diliana Pecheva, Pravesh Parekh, Diana M. Smith, Ashley Becker, Janosch Linkersdörfer, Richard Watts, Robert Loughnan, Donald J. Hagler, Kyung E. Rhee, Carolina Makowski, Anders M. Dale, Terry L. Jernigan

**Author notes:** Corresponding author: Alison Rigby, MS, MA, Neurosciences Graduate Program, University of California, San Diego, 9500 Gilman Dr., La Jolla, CA 92093.

## Abstract

**Importance:** Adolescent body mass index (BMI) is linked to brain microstructural alterations, but it is not known if this association is uniform across the BMI range.

**Objective:** To characterize the shape of the association between adolescent BMI and brain microstructure across the BMI range.

**Design:** Baseline, and 2-, 4-, and 6-year follow-up brain imaging and BMI from the Adolescent Brain Cognitive Development℠ Study, analyzed between June 2025 and June 2026.

**Setting:** Multicenter, population-based study including 21 sites distributed across the US.

**Participants:** 10,361 adolescents (5,034 [48.5%] female; aged 9-18 years) from this population-based cohort study were included after quality control of complete data.

**Exposures:** Centers for Disease Control BMI extended percentiles and percent from median BMI.

**Main Outcome and Measure:** The primary outcome was the restricted normalized isotropic (RNI) signal fraction derived from diffusion imaging which indexes water confined within spherical cellular structures and is sensitive to microstructural properties including cellularity.

**Results:** A total of 10,361 participants (5,034 [48.5%] female; aged 9-18 years) contributed 21,629 imaging observations across four timepoints. Higher BMI percentile was associated with greater RNI in bilateral reward- and appetite-related subcortical gray matter and white matter tracts, with associations varying spatially within structures, and with generally greater magnitude in males. Across these structures, the association increased steeply above the 80th percentile. This acceleration reflected compression of the percentile scale at high BMI; when BMI was reparameterized as percent from median, the association with RNI was approximately linear.

**Conclusions and Relevance:** Our results suggest that BMI is associated with brain microstructure in a graded manner within white matter tracts and subcortical gray matter related to reward, executive function, and appetitive control. This indicates that the association between BMI and microstructure is not confined to weight extremes and should be accounted for when modeling brain-body-behavior relationships during adolescence.

**Key Points:** *Question:* Does the association between adolescent body mass index (BMI) and brain microstructure vary across the BMI ranges?

*Findings:* In this longitudinal study, adolescent BMI percentile and percent from median BMI were significantly associated with brain microstructure in subcortical structures and white matter tracts. The rate of change above the 80th BMI percentile was steep, although this reflected a compression of effects in the upper percentile range; percent from median revealed an approximately linear relationship between BMI and brain microstructure.

*Meaning:* BMI is linearly associated with brain microstructure in adolescence, irrespective of clinically defined weight classes.

## Introduction

Adolescence is marked by dramatic change in body growth and brain organization. Adolescents gain roughly a fifth of their adult height and half of their adult weight alongside extensive neural remodeling^1–5^, while the neuroendocrine changes of puberty reshape the relative proportions of lean muscle mass and adipose tissue in a sex-specific manner^6^. However, how these bodily changes relate to concurrent brain development is less well characterized, highlighting a gap with clinical stakes. Body mass index (BMI) is the primary tool clinicians use to index growth and screen for weight-related health risks^7,8^. Both BMI extremes carry distinct health risks: high BMI signals obesity, which is linked to cardiovascular disease, type 2 diabetes, cancer, and depression^9,10^, while low BMI can reflect undernutrition^11^ and eating disorders^12–14^. Because BMI can be a marker for health risk, and has been linked to differences in brain structure during a period of rapid brain development^15–21^, understanding how BMI relates to brain development across the full BMI range is an important step toward identifying potential weight-related neurodevelopmental risk.

Diffusion-weighted imaging (DWI), which is sensitive to the diffusion of water through tissue, provides a noninvasive window into brain microstructure. Prior work using diffusion tensor imaging (DTI) has linked higher BMI to altered white matter (WM) organization^16,22^; however, tensor metrics are most informative in white matter tracts and difficult to interpret in gray matter (GM)^23^, conflate multiple microstructural properties into single summary measures^24^, and fail to resolve crossing fibers^25^, limiting biological specificity. Restriction spectrum imaging (RSI), an advanced multi-shell diffusion model, provides greater microstructural resolution in WM and GM. RSI decomposes the diffusion signal into restricted (intracellular), hindered (extracellular), and free-water compartments^25–29^. Its restricted normalized isotropic signal fraction (RNI) indexes isotropically restricted water, attributable primarily to intracellular compartments, but also captures other sources of variability^27,29^ and noise^30^. Because eating behavior and energy balance are governed by appetite and reward circuitry^31–33^, microstructure within these regions offers a biologically specific link between BMI and brain.

Using RSI in the Adolescent Brain Cognitive Development (ABCD) Study, prior work has reported positive associations between BMI and RNI in subcortical reward-related structures^15,17–20^. Given that RNI reflects cellularity, these studies have interpreted the positive BMI-RNI relationship as putative neuroinflammation via glial activation and infiltration^34^, a mechanism that is supported by animal models of obesity^35–37^ and that potentially explains evidence of altered cognitive function at high BMI^38^.

However, this literature is limited in several ways. First, and most consequential for clinical interpretation, the shape of the BMI-microstructure association across the full BMI range has not been characterized: prior studies have either modeled BMI as a linear predictor or contrasted discrete weight-status categories (e.g., with vs. without obesity), without examining how the association may be distributed across the BMI range. Whether the association is uniform or variable across the BMI range is an important distinction that bears directly on the clinical populations to whom these findings apply.

Second, most studies draw on only one or two early ABCD imaging timepoints, constraining the developmental window under examination. Lastly, region-of-interest (ROI) approaches average the effect across an entire structure leading to less nuanced characterization of the differential associations of BMI across the brain.

To address these gaps, we leveraged the ABCD Study® 6.1 release^39^, which includes four imaging timepoints spanning ages 9 to 18 years, providing a substantially wider developmental window than prior work. Using voxelwise nonlinear models, we characterized the sex-stratified association between BMI and RNI across the range of Centers for Disease Control and Prevention (CDC)-referenced BMI percentiles, adjusting for nonlinear effects of age and pubertal development, along with genetic ancestry, sociodemographic factors, and scanner serial number and software. Of note, CDC BMI percentiles are compressed in the upper tail, such that children with large differences in BMI map to nearly identical high percentile values^40^. To address this, we additionally used age- and sex-specific percent from median BMI (%median), which scales more directly with BMI, allowing a nuanced characterization of the BMI associations with microstructure in the extremity of the BMI range. We hypothesized that higher BMI would be associated with greater RNI in subcortical appetitive and reward-related structures and asked whether this association is uniform or variable across the BMI range. By mapping where along the BMI range and where in the brain these associations emerge, we aim to clarify the developmental and health relevance of BMI for adolescent brain microstructure.

## Methods

### Sample

We leveraged data from the ABCD Study (release 6.1)^39^, a large-scale longitudinal study enrolling 11,880 children at ages 9-11 across 21 US sites^41^. The University of California, San Diego Institutional Review Board approved all study protocols; parent/caregiver consent and child assent were obtained. Further details are described elsewhere^41–44^. We used data from baseline and 2-, 4-, and 6-year imaging follow-up visits. Demographics of the sample are described in Table 1. The final analysis sample included 10,361 unique participants (see eMethods in Supplement for exclusion criteria). This study follows the Strengthening the Reporting of Observational Studies in Epidemiology (STROBE) reporting guideline.

**Table 1.** Demographic Data for All Observations, Stratified by Sex and Study Timepoint.

| Characteristic | Overall | Baseline | 2-Year FU | 4-Year FU | 6-Year FU |
| --- | --- | --- | --- | --- | --- |
| Female |  |  |  |  |  |
| **No. of observations, No. (%)<sup>a</sup>** | 10,373 (48.0%) | 4,370 (42.1%) | 2,898 (27.9%) | 1,943 (18.7%) | 1,162 (11.2%) |
| **Age, y** | 12.0 (2.2) | 10.0 (0.6) | 12.0 (0.6) | 14.1 (0.7) | 16.0 (0.7) |
| **BMI, kg/m<sup>2</sup>** | 20.6 (4.9) | 18.7 (3.8) | 20.7 (4.6) | 22.8 (5.2) | 23.6 (5.4) |
| **CDC extended BMI percentile<sup>b</sup>** | 62.4 (29.1) | 59.4 (30.2) | 63.0 (29.2) | 67.1 (26.9) | 64.5 (27.0) |
| **Percent from median BMI, age/sex** | 13.9 (24.9) | 11.1 (22.7) | 14.7 (25.5) | 18.0 (27.0) | 15.8 (26.4) |
| **Puberty** | 3.1 (1.1) | 2.2 (0.8) | 3.2 (0.7) | 4.1 (0.5) | 4.5 (0.4) |
| **CDC weight class, No. (%)<sup>b</sup>** |  |  |  |  |  |
| Underweight | 263 (2.5%) | 150 (3.4%) | 70 (2.4%) | 25 (1.3%) | 18 (1.5%) |
| Healthy weight | 6,898 (66.5%) | 2,946 (67.4%) | 1,908 (65.8%) | 1,246 (64.1%) | 798 (68.7%) |
| Overweight | 1,648 (15.9%) | 667 (15.3%) | 465 (16.0%) | 340 (17.5%) | 176 (15.1%) |
| Obesity | 1,564 (15.1%) | 607 (13.9%) | 455 (15.7%) | 332 (17.1%) | 170 (14.6%) |
| **Race, No. (%)<sup>c</sup>** |  |  |  |  |  |
| White | 7,096 (68.4%) | 2,926 (67.0%) | 1,999 (69.0%) | 1,313 (67.6%) | 858 (73.8%) |
| Black or African American | 1,473 (14.2%) | 672 (15.4%) | 393 (13.6%) | 284 (14.6%) | 124 (10.7%) |
| Asian | 207 (2.0%) | 93 (2.1%) | 54 (1.9%) | 41 (2.1%) | 19 (1.6%) |
| American Indian or Alaska Native | 49 (0.5%) | 22 (0.5%) | 13 (0.4%) | 9 (0.5%) | 5 (0.4%) |
| Native Hawaiian or Other Pacific Islander | 14 (0.1%) | 4 (0.1%) | 5 (0.2%) | 3 (0.2%) | 2 (0.2%) |
| More than 1 race | 1,148 (11.1%) | 460 (10.5%) | 337 (11.6%) | 226 (11.6%) | 125 (10.8%) |
| Other or unknown | 385 (3.7%) | 192 (4.4%) | 97 (3.3%) | 67 (3.4%) | 29 (2.5%) |
| Missing | 1 (0.0%) | 1 (0.0%) | 0 (0.0%) | 0 (0.0%) | 0 (0.0%) |
| **Ethnicity, No. (%)<sup>d</sup>** |  |  |  |  |  |
| Hispanic | 2,010 (19.4%) | 860 (19.7%) | 554 (19.1%) | 395 (20.3%) | 201 (17.3%) |
| Non-Hispanic | 8,345 (80.4%) | 3,496 (80.0%) | 2,341 (80.8%) | 1,547 (79.6%) | 961 (82.7%) |
| Missing | 18 (0.2%) | 14 (0.3%) | 3 (0.1%) | 1 (0.1%) | 0 (0.0%) |
| **Annual household income, No. (%)** |  |  |  |  |  |
| <\$25 000 | 1,014 (9.8%) | 531 (12.2%) | 252 (8.7%) | 165 (8.5%) | 66 (5.7%) |
| \$25 000-\$49 999 | 1,241 (12.0%) | 585 (13.4%) | 363 (12.5%) | 206 (10.6%) | 87 (7.5%) |
| \$50 000-\$74 999 | 1,207 (11.6%) | 527 (12.1%) | 374 (12.9%) | 198 (10.2%) | 108 (9.3%) |
| \$75 000-\$99 999 | 1,388 (13.4%) | 610 (14.0%) | 405 (14.0%) | 237 (12.2%) | 136 (11.7%) |
| \$100 000-\$199 999 | 3,328 (32.1%) | 1,280 (29.3%) | 927 (32.0%) | 679 (34.9%) | 442 (38.0%) |
| ≥\$200 000 | 1,452 (14.0%) | 492 (11.3%) | 380 (13.1%) | 322 (16.6%) | 258 (22.2%) |
| Declined to answer | 369 (3.6%) | 174 (4.0%) | 83 (2.9%) | 74 (3.8%) | 38 (3.3%) |
| Don't know | 374 (3.6%) | 171 (3.9%) | 114 (3.9%) | 62 (3.2%) | 27 (2.3%) |
| **Highest parental education, No. (%)** |  |  |  |  |  |
| Up to high school (no diploma) | 478 (4.6%) | 224 (5.1%) | 120 (4.1%) | 91 (4.7%) | 43 (3.7%) |
| High school diploma/GED | 825 (8.0%) | 361 (8.3%) | 224 (7.7%) | 152 (7.8%) | 88 (7.6%) |

| Characteristic (continued) | Overall | Baseline | 2-Year FU | 4-Year FU | 6-Year FU |
| --- | --- | --- | --- | --- | --- |
| **Highest parental education, No. (%)** |  |  |  |  |  |
| Some college | 2,839 (27.4%) | 1,071 (24.5%) | 902 (31.1%) | 547 (28.2%) | 319 (27.5%) |
| Bachelor's degree | 2,465 (23.8%) | 1,127 (25.8%) | 636 (21.9%) | 435 (22.4%) | 267 (23.0%) |
| Graduate or professional degree | 3,766 (36.3%) | 1,587 (36.3%) | 1,016 (35.1%) | 718 (37.0%) | 445 (38.3%) |
| Male |  |  |  |  |  |
| **No. of observations, No. (%)<sup>a</sup>** | 11,256 (52.0%) | 4,118 (36.6%) | 3,001 (26.7%) | 2,591 (23.0%) | 1,546 (13.7%) |
| **Age, y** | 12.3 (2.3) | 10.0 (0.6) | 12.1 (0.7) | 14.2 (0.7) | 16.1 (0.7) |
| **BMI, kg/m<sup>2</sup>** | 20.5 (4.7) | 18.5 (3.6) | 20.2 (4.3) | 22.2 (5.0) | 23.0 (4.8) |
| **CDC extended BMI percentile<sup>b</sup>** | 61.9 (30.1) | 60.9 (29.9) | 62.2 (30.4) | 63.9 (30.0) | 60.5 (30.2) |
| **Percent from median BMI, age/sex** | 13.0 (23.7) | 11.3 (21.6) | 13.5 (24.1) | 15.6 (26.2) | 11.8 (23.4) |
| **Puberty** | 2.4 (1.1) | 1.5 (0.5) | 2.2 (0.7) | 3.2 (0.7) | 3.8 (0.4) |
| **CDC weight class, No. (%)<sup>b</sup>** |  |  |  |  |  |
| Underweight | 355 (3.2%) | 107 (2.6%) | 105 (3.5%) | 85 (3.3%) | 58 (3.8%) |
| Healthy weight | 7,281 (64.7%) | 2,762 (67.1%) | 1,900 (63.3%) | 1,591 (61.4%) | 1,028 (66.5%) |
| Overweight | 1,715 (15.2%) | 608 (14.8%) | 489 (16.3%) | 401 (15.5%) | 217 (14.0%) |
| Obesity | 1,905 (16.9%) | 641 (15.6%) | 507 (16.9%) | 514 (19.8%) | 243 (15.7%) |
| **Race, No. (%)<sup>c</sup>** |  |  |  |  |  |
| White | 7,994 (71.0%) | 2,860 (69.5%) | 2,145 (71.5%) | 1,823 (70.4%) | 1,166 (75.4%) |
| Black or African American | 1,436 (12.8%) | 568 (13.8%) | 376 (12.5%) | 338 (13.0%) | 154 (10.0%) |
| Asian | 223 (2.0%) | 84 (2.0%) | 54 (1.8%) | 50 (1.9%) | 35 (2.3%) |
| American Indian or Alaska Native | 41 (0.4%) | 11 (0.3%) | 12 (0.4%) | 11 (0.4%) | 7 (0.5%) |
| Native Hawaiian or Other Pacific Islander | 22 (0.2%) | 7 (0.2%) | 7 (0.2%) | 7 (0.3%) | 1 (0.1%) |
| More than 1 race | 1,183 (10.5%) | 430 (10.4%) | 313 (10.4%) | 291 (11.2%) | 149 (9.6%) |
| Other or unknown | 357 (3.2%) | 158 (3.8%) | 94 (3.1%) | 71 (2.7%) | 34 (2.2%) |
| Missing | 0 (0.0%) | 0 (0.0%) | 0 (0.0%) | 0 (0.0%) | 0 (0.0%) |
| **Ethnicity, No. (%)<sup>d</sup>** |  |  |  |  |  |
| Hispanic | 2,156 (19.2%) | 834 (20.3%) | 567 (18.9%) | 483 (18.6%) | 272 (17.6%) |
| Non-Hispanic | 9,060 (80.5%) | 3,261 (79.2%) | 2,424 (80.8%) | 2,103 (81.2%) | 1,272 (82.3%) |
| Missing | 40 (0.4%) | 23 (0.6%) | 10 (0.3%) | 5 (0.2%) | 2 (0.1%) |
| **Annual household income, No. (%)** |  |  |  |  |  |
| <\$25 000 | 1,114 (9.9%) | 493 (12.0%) | 297 (9.9%) | 220 (8.5%) | 104 (6.7%) |
| \$25 000-\$49 999 | 1,222 (10.9%) | 514 (12.5%) | 314 (10.5%) | 264 (10.2%) | 130 (8.4%) |
| \$50 000-\$74 999 | 1,329 (11.8%) | 528 (12.8%) | 393 (13.1%) | 274 (10.6%) | 134 (8.7%) |
| \$75 000-\$99 999 | 1,440 (12.8%) | 558 (13.6%) | 399 (13.3%) | 315 (12.2%) | 168 (10.9%) |
| \$100 000-\$199 999 | 3,626 (32.2%) | 1,220 (29.6%) | 987 (32.9%) | 851 (32.8%) | 568 (36.7%) |
| ≥\$200 000 | 1,657 (14.7%) | 462 (11.2%) | 391 (13.0%) | 457 (17.6%) | 347 (22.4%) |
| **Annual household income, No. (%)** |  |  |  |  |  |
| Declined to answer | 468 (4.2%) | 185 (4.5%) | 108 (3.6%) | 114 (4.4%) | 61 (3.9%) |
| Don't know | 400 (3.6%) | 158 (3.8%) | 112 (3.7%) | 96 (3.7%) | 34 (2.2%) |
| **Highest parental education, No. (%)** |  |  |  |  |  |
| Up to high school (no diploma) | 405 (3.6%) | 150 (3.6%) | 99 (3.3%) | 96 (3.7%) | 60 (3.9%) |
| High school diploma/GED | 929 (8.3%) | 353 (8.6%) | 253 (8.4%) | 217 (8.4%) | 106 (6.9%) |
| Some college | 3,231 (28.7%) | 1,087 (26.4%) | 947 (31.6%) | 770 (29.7%) | 427 (27.6%) |
| Bachelor's degree | 2,666 (23.7%) | 1,073 (26.1%) | 688 (22.9%) | 565 (21.8%) | 340 (22.0%) |
| Graduate or professional degree | 4,025 (35.8%) | 1,455 (35.3%) | 1,014 (33.8%) | 943 (36.4%) | 613 (39.7%) |

Abbreviations: BMI, body mass index; CDC, Centers for Disease Control and Prevention; FU, Follow-up; GED, Generalized Education Degree; SD, standard deviation.
<sup>a</sup> Categorical values are presented as No. (%), with percentages computed within each column of the corresponding sex block; continuous variables are presented as mean (SD). For the number of observations row, percentages in the Overall column are based on all observations in the analytic sample; percentages in study wave columns are based on all observations within each sex.
<sup>b</sup> CDC extended BMI percentile was derived using CDC age- and sex-specific growth reference data. CDC weight classes were derived from CDC extended BMI percentile as underweight (<5th percentile), healthy weight (5th to <85th percentile), overweight (85th to <95th percentile), and obesity (≥95th percentile).
<sup>c</sup> Race (NIH 7 categories) was self-reported by the caregiver. Descriptive statistics are reported for self-reported race only for interpretation of sample diversity.
<sup>d</sup> Ethnicity (2 categories) was self-reported by the caregiver.
<sup>e</sup> A total of 21,629 observations were included (10,373 female; 11,256 male). The sample comprised 10,361 unique participants (5,034 female; 5,327 male). Observations by study timepoint (all participants): Baseline: 8,488; 2-Year FU: 5,899; 4-Year FU: 4,534; 6-Year FU: 2,708. Characteristics are summarized across all eligible imaging visits in the voxelwise model sample. Female and male results are presented in stacked blocks. Within each block, the Overall column summarizes all observations for that sex, and wave
columns summarize observations within that sex and study wave. Column percentages are computed within each column of the corresponding sex block.

**Table 2.**
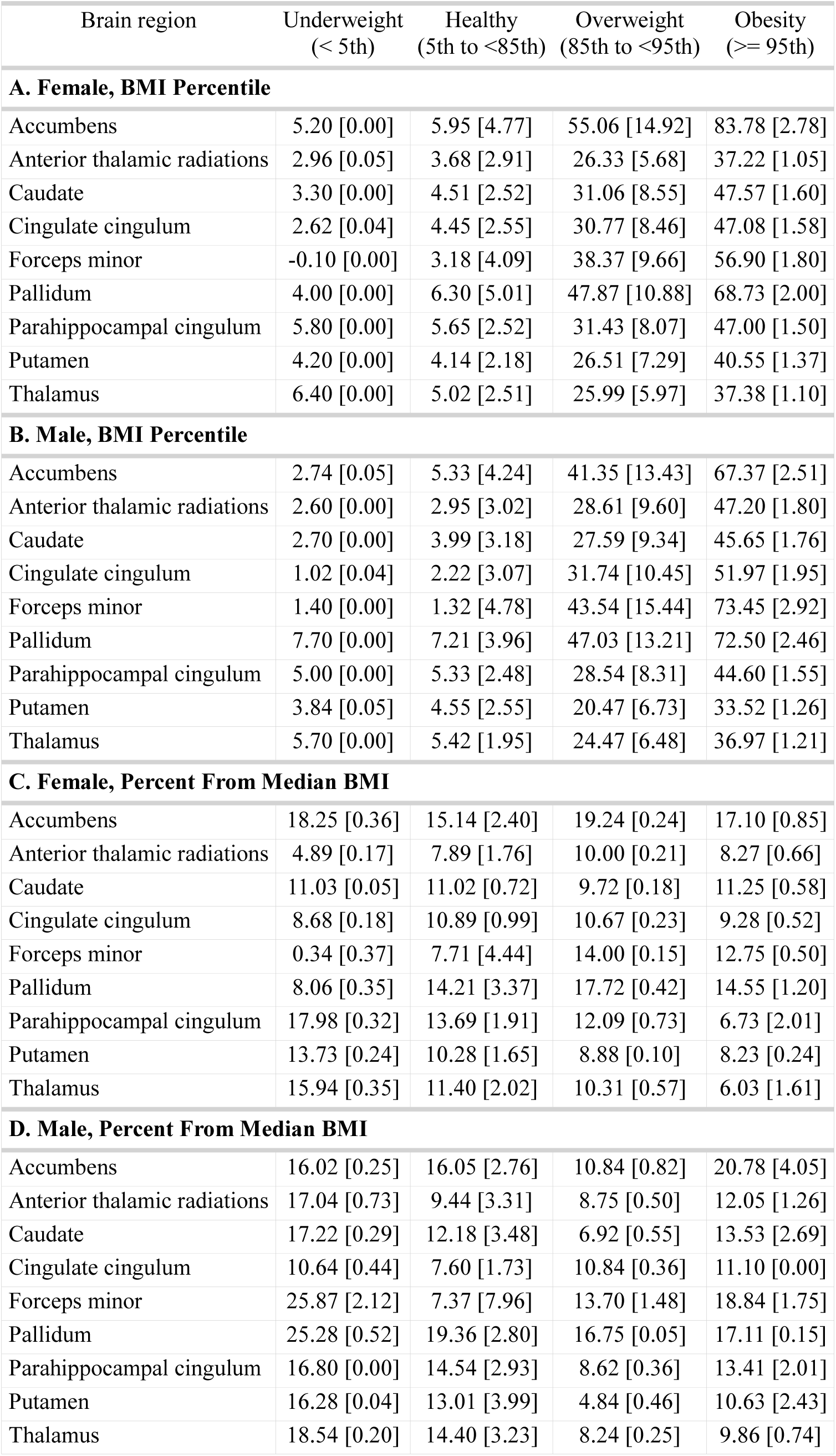
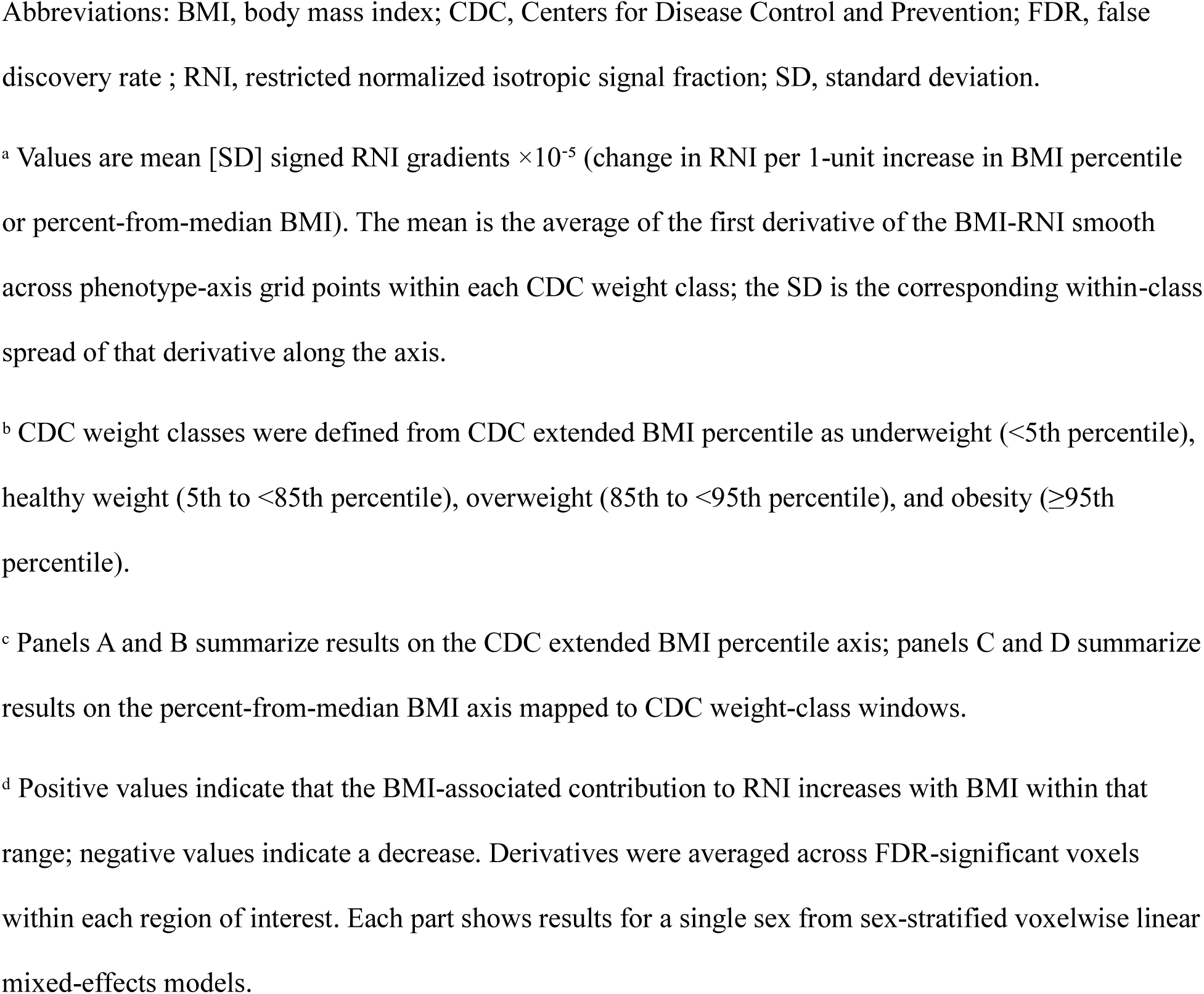
Mean Rate of Change in the BMI-RNI Association by CDC Weight Class, Stratified by Sex^a,b,c,d^.

| Brain region | Underweight<br>(< 5th) | Healthy<br>(5th to <85th) | Overweight<br>(85th to <95th) | Obesity<br>(≥ 95th) |
| --- | --- | --- | --- | --- |
| **A. Female, BMI Percentile** |  |  |  |  |
| Accumbens | 5.20 [0.00] | 5.95 [4.77] | 55.06 [14.92] | 83.78 [2.78] |
| Anterior thalamic radiations | 2.96 [0.05] | 3.68 [2.91] | 26.33 [5.68] | 37.22 [1.05] |
| Caudate | 3.30 [0.00] | 4.51 [2.52] | 31.06 [8.55] | 47.57 [1.60] |
| Cingulate cingulum | 2.62 [0.04] | 4.45 [2.55] | 30.77 [8.46] | 47.08 [1.58] |
| Forceps minor | -0.10 [0.00] | 3.18 [4.09] | 38.37 [9.66] | 56.90 [1.80] |
| Pallidum | 4.00 [0.00] | 6.30 [5.01] | 47.87 [10.88] | 68.73 [2.00] |
| Parahippocampal cingulum | 5.80 [0.00] | 5.65 [2.52] | 31.43 [8.07] | 47.00 [1.50] |
| Putamen | 4.20 [0.00] | 4.14 [2.18] | 26.51 [7.29] | 40.55 [1.37] |
| Thalamus | 6.40 [0.00] | 5.02 [2.51] | 25.99 [5.97] | 37.38 [1.10] |
| **B. Male, BMI Percentile** |  |  |  |  |
| Accumbens | 2.74 [0.05] | 5.33 [4.24] | 41.35 [13.43] | 67.37 [2.51] |
| Anterior thalamic radiations | 2.60 [0.00] | 2.95 [3.02] | 28.61 [9.60] | 47.20 [1.80] |
| Caudate | 2.70 [0.00] | 3.99 [3.18] | 27.59 [9.34] | 45.65 [1.76] |
| Cingulate cingulum | 1.02 [0.04] | 2.22 [3.07] | 31.74 [10.45] | 51.97 [1.95] |
| Forceps minor | 1.40 [0.00] | 1.32 [4.78] | 43.54 [15.44] | 73.45 [2.92] |
| Pallidum | 7.70 [0.00] | 7.21 [3.96] | 47.03 [13.21] | 72.50 [2.46] |
| Parahippocampal cingulum | 5.00 [0.00] | 5.33 [2.48] | 28.54 [8.31] | 44.60 [1.55] |
| Putamen | 3.84 [0.05] | 4.55 [2.55] | 20.47 [6.73] | 33.52 [1.26] |
| Thalamus | 5.70 [0.00] | 5.42 [1.95] | 24.47 [6.48] | 36.97 [1.21] |
| **C. Female, Percent From Median BMI** |  |  |  |  |
| Accumbens | 18.25 [0.36] | 15.14 [2.40] | 19.24 [0.24] | 17.10 [0.85] |
| Anterior thalamic radiations | 4.89 [0.17] | 7.89 [1.76] | 10.00 [0.21] | 8.27 [0.66] |
| Caudate | 11.03 [0.05] | 11.02 [0.72] | 9.72 [0.18] | 11.25 [0.58] |
| Cingulate cingulum | 8.68 [0.18] | 10.89 [0.99] | 10.67 [0.23] | 9.28 [0.52] |
| Forceps minor | 0.34 [0.37] | 7.71 [4.44] | 14.00 [0.15] | 12.75 [0.50] |
| Pallidum | 8.06 [0.35] | 14.21 [3.37] | 17.72 [0.42] | 14.55 [1.20] |
| Parahippocampal cingulum | 17.98 [0.32] | 13.69 [1.91] | 12.09 [0.73] | 6.73 [2.01] |
| Putamen | 13.73 [0.24] | 10.28 [1.65] | 8.88 [0.10] | 8.23 [0.24] |
| Thalamus | 15.94 [0.35] | 11.40 [2.02] | 10.31 [0.57] | 6.03 [1.61] |
| **D. Male, Percent From Median BMI** |  |  |  |  |
| Accumbens | 16.02 [0.25] | 16.05 [2.76] | 10.84 [0.82] | 20.78 [4.05] |
| Anterior thalamic radiations | 17.04 [0.73] | 9.44 [3.31] | 8.75 [0.50] | 12.05 [1.26] |
| Caudate | 17.22 [0.29] | 12.18 [3.48] | 6.92 [0.55] | 13.53 [2.69] |
| Cingulate cingulum | 10.64 [0.44] | 7.60 [1.73] | 10.84 [0.36] | 11.10 [0.00] |
| Forceps minor | 25.87 [2.12] | 7.37 [7.96] | 13.70 [1.48] | 18.84 [1.75] |
| Pallidum | 25.28 [0.52] | 19.36 [2.80] | 16.75 [0.05] | 17.11 [0.15] |
| Parahippocampal cingulum | 16.80 [0.00] | 14.54 [2.93] | 8.62 [0.36] | 13.41 [2.01] |
| Putamen | 16.28 [0.04] | 13.01 [3.99] | 4.84 [0.46] | 10.63 [2.43] |
| Thalamus | 18.54 [0.20] | 14.40 [3.23] | 8.24 [0.25] | 9.86 [0.74] |

Abbreviations: BMI, body mass index; CDC, Centers for Disease Control and Prevention; FDR, false discovery rate ; RNI, restricted normalized isotropic signal fraction; SD, standard deviation.
<sup>a</sup> Values are mean [SD] signed RNI gradients $\times 10^{-5}$ (change in RNI per 1-unit increase in BMI percentile or percent-from-median BMI). The mean is the average of the first derivative of the BMI-RNI smooth across phenotype-axis grid points within each CDC weight class; the SD is the corresponding within-class spread of that derivative along the axis.
<sup>b</sup> CDC weight classes were defined from CDC extended BMI percentile as underweight (<5th percentile), healthy weight (5th to <85th percentile), overweight (85th to <95th percentile), and obesity ( $\geq 95$ th percentile).
<sup>c</sup> Panels A and B summarize results on the CDC extended BMI percentile axis; panels C and D summarize results on the percent-from-median BMI axis mapped to CDC weight-class windows.
<sup>d</sup> Positive values indicate that the BMI-associated contribution to RNI increases with BMI within that range; negative values indicate a decrease. Derivatives were averaged across FDR-significant voxels within each region of interest. Each part shows results for a single sex from sex-stratified voxelwise linear mixed-effects models.

### BMI Phenotypes

We computed BMI as weight(kg) / height(m)^2^ and calculated the CDC age- and sex-specific extended BMI percentiles and the %median using the *cdcanthro* package in R^45–47^. To illustrate the relationship between these two BMI metrics, we plotted each imaging visit’s CDC extended BMI percentile against its corresponding %median, stratified according to sex with CDC weight class-colored mappings (Figure 1).

**Figure 1.**
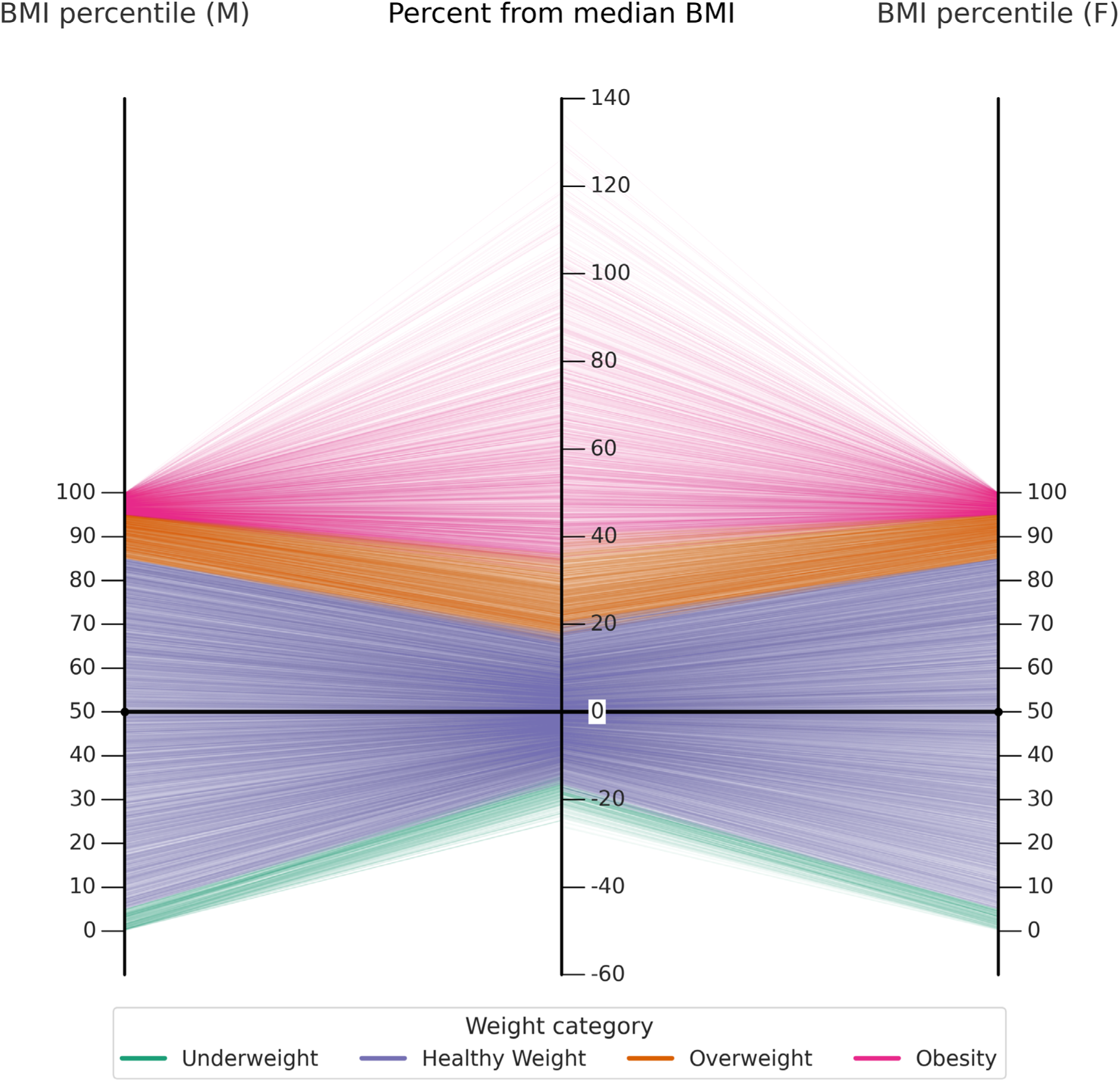
Mapping of CDC extended BMI percentile to percent-from-median BMI. Each line connects an individual’s age-and sex-specific BMI percentile at a given timepoint (left and right axes correspond to males and females, respectively) to its percent deviation from age-and sex-specific median BMI (center axis), colored by CDC weight class.

### Neuroimaging Acquisition and Processing

Details on the ABCD Study imaging acquisition, processing, and atlas registration protocol^48,49^, and descriptions of the RSI model derivation and its application to the ABCD Study data^25,26^ can be found elsewhere. From the ABCD Study concatenated voxelwise RSI imaging data^50^, we used the RNI diffusion signal fraction for our analyses across the whole brain.

### Statistical Analysis

We used voxelwise, sex-stratified, univariate generalized additive mixed-effects models to estimate associations between BMI phenotypes and RNI. Given the sex-specific effects of puberty^51,52^, we stratified models by sex. We examined BMI-RSI associations using the following model:

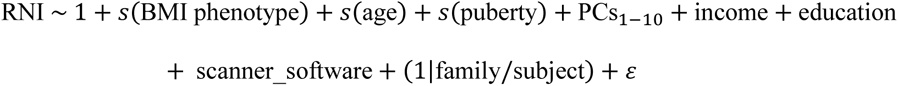

where, generically *s* denotes a smooth function and (1|family/subject) denotes random intercepts for repeated measurements of subjects nested within families. We constructed nonlinear terms for BMI phenotype, age, and puberty using natural cubic splines with unit heights at knots^53,54^ (further details in eMethods in Supplement). We included household income and highest parental education as categorical covariates, the first 10 genetic principal components to account for population stratification, and dummy-coded scanner_software to account for MRI hardware differences. We used the Fast and Efficient Mixed-Effects Analysis (FEMA) toolbox v4.0.3^54,55^ to estimate voxelwise regression coefficients. To test the association between BMI phenotype and RNI at each voxel, we performed an omnibus Wald test on the joint null that all BMI spline coefficients were zero (no association). We corrected voxelwise results for multiple comparisons using the Benjamini-Hochberg false discovery rate (FDR) procedure at *q* < 0.05, applied under the positive dependency assumption^56^. We compared the sex-stratified maps using Pearson correlation of unthresholded spline Wald statistics and Dice similarity of voxels surviving FDR. Analyses used R version 4.5.3^57^, MATLAB R2022a^58^, and Python 3.13.13^59^.

To visualize the shape of BMI-RNI associations for regions with the most significant voxelwise associations, we conducted follow-up analyses in nine regions of interest, selected as those with the highest-ranking omnibus Wald statistics across both males and females, including the nucleus accumbens, caudate, pallidum, putamen, and thalamus (FreeSurfer aseg segmentation^60^) and the forceps minor, anterior thalamic radiations, cingulate cingulum, and parahippocampal cingulum (AtlasTrack^61^). Details on voxel extraction, reconstruction of the BMI-RNI smooth function, and calculation of its first derivatives, or gradients (the change in RNI per BMI unit) are provided in eMethods in Supplement.

### Sensitivity Analysis

Given the recently reported association between acquired flip angle and body size in dMRI data acquired on Siemens VE11 with the MGH sequence^62^, we performed a sensitivity analysis comparing two subsets of participants: observations with data acquired on VE11 and observations with data acquired on all other scanner software versions (see eMethods in Supplement).

## Results

A total of 10,361 unique participants (5,034 [48.5%] female) with 21,629 observations (10,373 [48.0%] female) across the four neuroimaging timepoints were included in the analyses.

### Spatial distribution of BMI percentile associations with subcortical GM and WM tract microstructure

We observed significant positive associations of the smooth function of BMI percentile with RNI bilaterally in the following subcortical GM structures: nucleus accumbens, caudate, pallidum, putamen, and thalamus and the following WM tracts: forceps minor, cingulate cingulum, and parahippocampal cingulum. The strength of significance varied in spatial distribution within structures, shown in Figure 2. For example, the medial posterior portion of the forceps minor, dorsal portions of the pallidum, anterior portions of the putamen, and posterior portions of the thalamus were most significantly associated with BMI percentile relative to the rest of each structure. This spatial pattern was similar between sexes (eTable 2 in Supplement).

**Figure 2.**
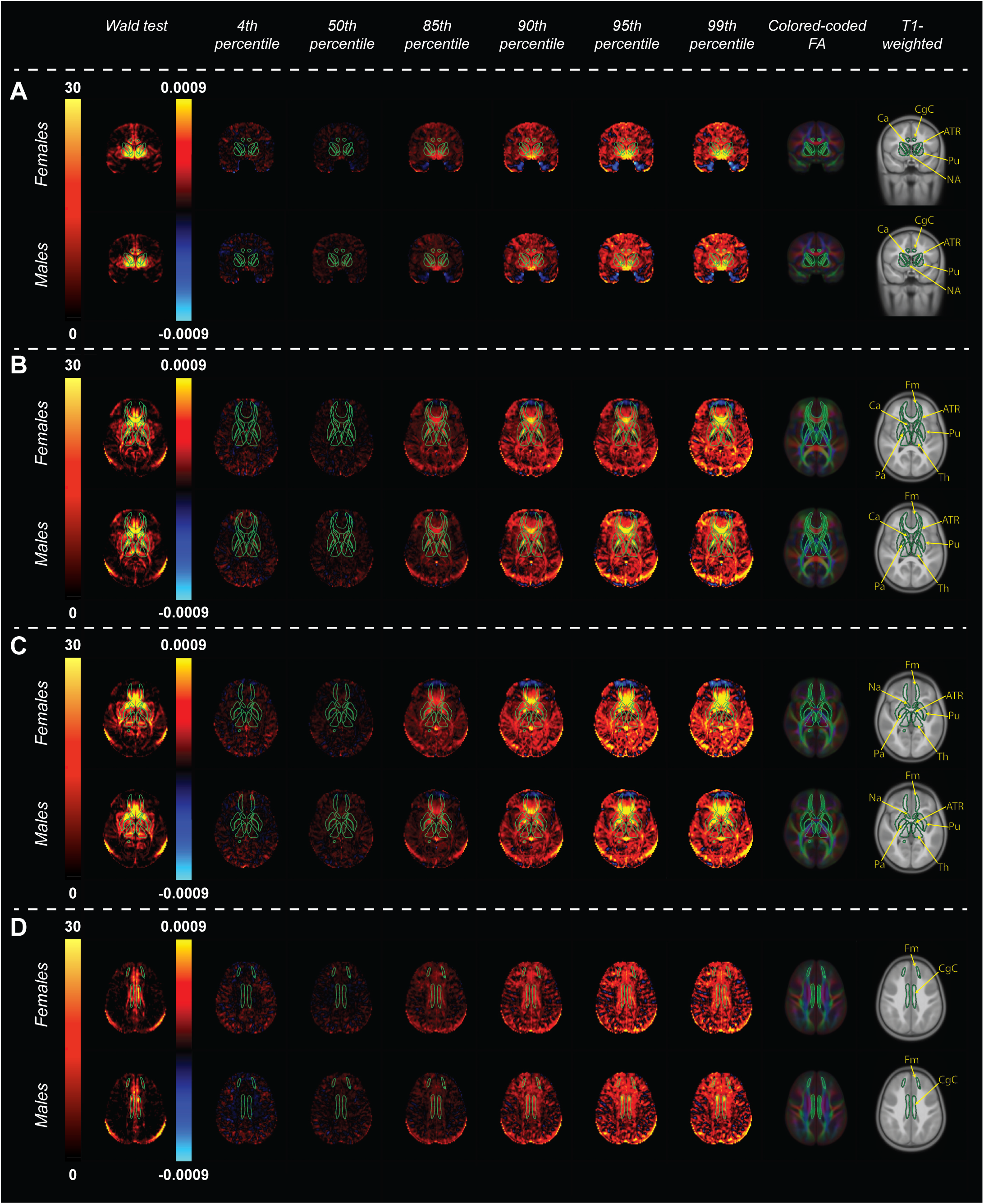
Voxelwise BMI-RNI associations and local rates of change across the BMI percentile range. Column 1. Voxelwise -log₁₀ *p*-value maps from the omnibus Wald test of the nonlinear BMI percentile effect on restricted normalized isotropic signal fraction (RNI), thresholded at FDR q < 0.05. Columns 2-7. Derivative maps of the BMI-RNI smooth, showing the estimated instantaneous rate of change in the BMI effect on RNI per unit BMI percentile at the 4^th^ (underweight), 50^th^ (healthy weight), 85^th^ & 90^th^ (overweight), and 95^th^ & 99^th^ (obesity) BMI percentiles. Warm colors indicate that the BMI-associated contribution to RNI increased with BMI percentile at that location; cool colors indicate a decrease. Columns 8-9. Color-coded fractional anisotropy (left-right: red, anterior-posterior: green, inferior-superior: blue) and T1-weighted images for anatomical reference. **(A)** Coronal view highlighting the nucleus accumbens, anterior thalamic radiations, caudate, and putamen. **(B)** Axial view highlighting the anterior thalamic radiations, caudate, forceps minor, pallidum, putamen, and thalamus**. (C)** Axial view highlighting the accumbens, anterior thalamic radiations, forceps minor, pallidum, putamen, and thalamus. **(D)** Axial view highlighting the cingulate cingulum. Subcortical ROI outlines and labels are shown from the FreeSurfer automated subcortical segmentation and AtlasTrack. All maps were derived from sex-stratified voxelwise linear mixed-effects models including smooth terms for BMI percentile, age, and pubertal development, with covariates for genetic principal components, household income, highest parental education, and scanner/software version. The bright band at the posterior of the brain on axial slices is a chemical-shift (fat) artifact and does not reflect brain microstructure. Label abbreviations: NA = nucleus accumbens, ATR = anterior thalamic radiations; Ca = caudate; CgC = cingulate cingulum; Fm = forceps minor; Pa = pallidum; Pu = putamen; Th = thalamus.

Because BMI was modeled as a smooth function, the association has no single slope; we therefore characterized it by the first derivative of the fitted smooth, or the rate of change in estimated RNI per unit BMI percentile at each point along the range. Voxelwise derivative maps corresponding to BMI percentiles at the CDC-defined weight classes -4^th^ (underweight), 50^th^ (healthy), 85^th^ & 90^th^ (overweight), and 95^th^ & 99^th^ (obesity) percentiles -are shown in Figure 2 and in movies spanning the full percentile range in eVideos 1-8 in Supplement. The derivative was positive across regions, and its magnitude substantially increased above the 85th percentile (overweight threshold) and was steepest at the 99th percentile in all nine structures, indicating that the association of BMI with RNI rose most steeply at the highest BMI percentiles.

### Nonlinear pattern of BMI percentile-RNI associations across the BMI percentile distribution

Figure 3A depicts the estimated associations between BMI percentiles and RNI across the nine structures. The estimated associations between BMI and RNI increased monotonically with BMI percentile, rising subtly across lower and middle percentiles before increasing steeply above the 80th. While the overall magnitude of the association was consistently higher for males than females across most structures, sex differences were most pronounced in the nucleus accumbens, anterior thalamic radiations, and caudate, where confidence intervals showed minimal overlap across the BMI percentile range. In contrast, male and female effects were largely overlapping in the cingulum and putamen.

**Figure 3.**
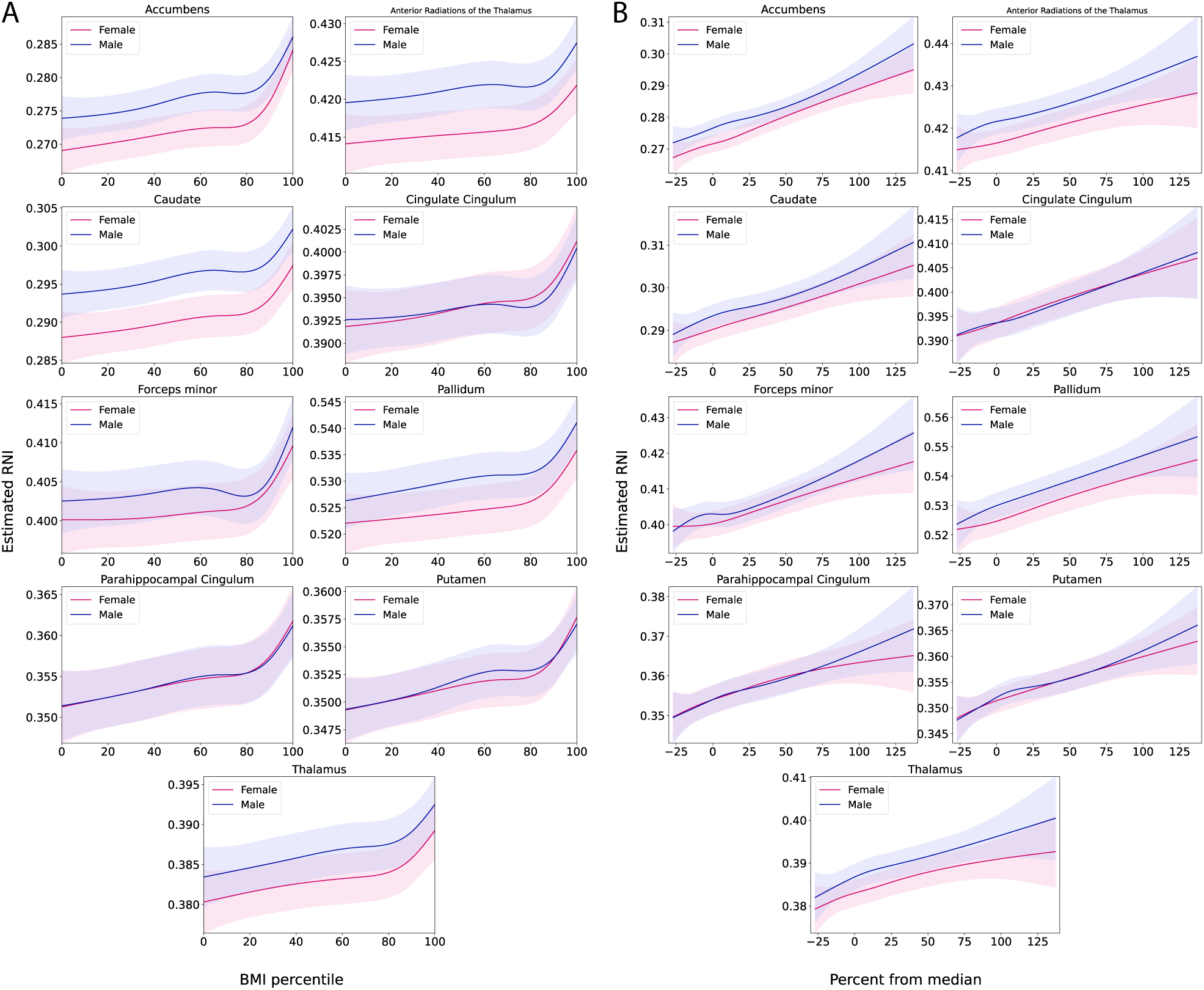
Sex-stratified BMI-RNI associations in selected regions of interest. Curves show the region-averaged contribution of the BMI spline (plus intercept) on the restricted normalized isotropic signal fraction (RNI) as a function of **(A)** CDC extended BMI percentile and **(B)** percent from median BMI, among voxels with an FDR-significant BMI-RNI association. Shaded bands indicate averaged voxelwise 95% confidence intervals. Models were estimated separately in females and males.

Notably, despite the higher overall magnitude for males, females showed steeper rates of change in the nucleus accumbens above the 80th percentile (Figure 3).

We quantified the mean rate of change within CDC weight classes according to BMI percentiles for females and males, respectively (Tables 2A and 2B). For example, in the nucleus accumbens bilaterally, the mean rate of change in RNI per BMI percentile was 14.1-fold greater for obesity (>=95^th^ percentile) and 9.3-fold for overweight (85^th^ to <95^th^ percentile) than for healthy weight categories (5^th^ to <85^th^ percentile) for females, and 12.6-fold and 7.8-fold greater, respectively, in males for the same weight categories. Across subcortical structures, the rates of change were consistently several-fold greater for obesity and overweight than for healthy weight and underweight (<5^th^ percentile) classes, ranging from 4.1-fold between overweight and underweight categories in the thalamus for females, to 55.6-fold difference between obesity and healthy weight in the forceps minor for males.

### Linear relationship between percent from median BMI and RNI

The steepening of the BMI percentile-RNI association above the 80^th^ percentile was striking. To evaluate whether it reflected compression of CDC BMI percentile at its upper extreme (Figure 1), we reparameterized BMI as %median, which is not compressed at extremes and scales more linearly with BMI^45,63^. This range is well-represented in ABCD: 37.6% of the sample fell above the 80th percentile, allowing us to characterize the upper BMI range in depth. Under this parameterization, the relationship between %median and RNI was approximately linear, as visualized in the nine structures that showed notable BMI associations in the voxelwise maps (Figure 3B, Tables 2C and 2D, and eFigure 9 and eVideos 9-16 in Supplement). Consistent with an approximately linear association, the rate of change in RNI with %median was relatively stable across weight classes and was comparable between females and males (e.g., nucleus accumbens mean gradient ≈ 17.4 vs. 15.9 ×10⁻⁵), with the steepest slopes in the nucleus accumbens (females) and pallidum (males). The spatial distribution mirrored the percentile results, remaining similar between sexes (eTable 2 in Supplement).

### Sensitivity analysis: Similar BMI-RNI associations in VE11 and non-VE11 samples

Both VE11 and non-VE11 groups showed similar spatial distribution and effect sizes of BMI phenotype-RNI associations compared with the main results (see eResults, eFigures 10-14, and eTable 1 in Supplement).

## Discussion

In this longitudinal study of 10,361 adolescents ages 9-18 years, higher BMI was associated with greater RNI (a diffusion MRI marker of tissue cellularity) in reward-related brain circuitry in a graded manner across the full BMI range, rather than emerging at clinical obesity. These associations spanned subcortical reward-related structures, prefrontal WM tracts, and limbic tracts. An apparent threshold above the 80th percentile reflected percentile scale compression; under an uncompressed metric (%median), the association between BMI and brain microstructure was graded and linear across the full BMI range rather than dynamic at BMI extremes. In other words, the shape of the association depended on the chosen scale.

Prior ABCD studies have consistently reported positive linear associations between BMI and microstructure in WM tracts and subcortical GM involved in reward, executive function, and appetitive control^15,17–19,22,64,65^. Our graded finding is therefore consistent with, rather than contradictory to, prior work - a linear model fit to a monotonic, threshold-free association will still detect a positive linear association, exactly what those studies reported. Our study extends this literature by characterizing the shape of the association directly rather than assuming it, aided by the wider developmental window of the ABCD 6.1 release (four imaging timepoints spanning ages 9-18), voxelwise resolution, and the parameterization of BMI as %median.

Because RNI indexes isotropically restricted intracellular water, it is interpreted as a marker of cellularity or tissue density rather than of any single cell type^27,29^. Prior work has proposed that greater RNI in relation to higher BMI reflects neuroinflammation^34^, namely glial activation and infiltration, a mechanism supported by animal models of obesity^35–37^ and a process to which dMRI is sensitive^66^.

However, this signal is not specific to any one process: glia that mediate inflammatory responses also carry out the synaptic pruning and circuit refinement^67–72^ that characterize normative adolescent brain and physical development. Greater RNI in relation to BMI could therefore reflect inflammatory processes or the typical neural remodeling that accompanies physical maturation, and current diffusion measures cannot distinguish the two; approaches with greater cell-type specificity, such as diffusion signatures tuned to glial morphology^66^, will be needed to adjudicate.

The anatomical distribution of our findings underscores this ambiguity. In animal models, diet-induced neuroinflammation arises primarily in the hypothalamus^35–37^, which is the principal regulator of energy homeostasis and weight stability. Yet in our analyses the hypothalamus -the canonical homeostatic and putative neuroinflammatory target -was not among the structures with the most highly significant BMI associations, which were instead concentrated primarily in reward-related regions and their associated white matter connections that link subcortical, prefrontal, and limbic areas^32^. The prominence of associations in striatal and reward-related circuitry aligns with models positing that dysregulation of dopaminergic reward pathways is central, rather than secondary, in obesity^73,74^. Our findings may reflect a heightened involvement of reward compared to homeostatic circuitry in the BMI-microstructure relationship that is specific to the adolescent developmental window and merits further study.

While our findings confirm that modeling BMI as a linear predictor is sufficient, we encourage the use of nonlinear approaches to examine the shape of relationships between variables before assuming linearity. Importantly, given its widespread associations with brain microstructure, BMI should be considered as a covariate when modeling brain-body-behavior relationships in adolescence, where relevant to the research question. Beyond developmental research, our findings also carry a direct clinical implication: if the association between BMI and brain microstructure is largely linear, there is no weight class threshold below which BMI is unrelated to the developing brain, prompting consideration of how BMI is clinically scaled. BMI percentile, the metric clinicians use to chart growth and classify weight status, compresses a broad range of BMI into a narrow percentile band^40^. Because the BMI-microstructure association we observed is graded and continuous, a scale with limited resolution at the high end can distort the apparent shape of that association. As a clinical scale, BMI percentiles may be limited in assessing clinically-relevant nuances at the upper extremes.

### Strengths and Limitations

Our study’s strengths include its wide developmental window, voxelwise resolution, and direct modeling of the association’s shape. However, we note that BMI is an indirect measure of body composition and adiposity. Moreover, its relationship with health risk varies across genetic ancestry, limiting its interpretability across populations^75,76^. While we included puberty as a covariate, the ABCD pubertal measures may not accurately capture pubertal processes as they are based on self-reports and are relatively noisy. Diffusion metrics, including RNI, cannot index specific cell types, and therefore, we are limited in mechanistic interpretation.

For larger-bodied participants, a reduced acquired flip angle was identified in the dMRI data acquired on Siemens scanners running the MGH sequence on VE11 software^62^. This resulted in reduced signal-to-noise ratio, potentially confounding derived dMRI metrics, thereby affecting the association between BMI and brain microstructure. Our sensitivity analysis showed that BMI-RNI associations were present in non-VE11 data, similar to our primary analyses and VE11-only data, indicating that the associations are not driven by this potential confound. However, further study is required to fully characterize these effects.

## Conclusions

The shape of the association between BMI and microstructure in WM tracts and subcortical GM related to reward, executive function, and appetitive control is mostly linear across adolescence, rather than dynamic at clinically-relevant thresholds. Given the positive associations between BMI with brain microstructure during development, researchers should strongly consider incorporating BMI as a covariate in brain-body-behavior models, but use age-and sex-adjusted percent from median BMI rather than BMI percentile.

## Acknowledgements

Data used in the preparation of this article were obtained from the Adolescent Brain Cognitive Development™ (ABCD) Study, held in the NIH Brain Development Cohorts Data Sharing Platform. This is a multisite, longitudinal study designed to recruit more than 10,000 children aged 9–10 and follow them over 10 years into early adulthood. The ABCD Study® is supported by the National Institutes of Health and additional federal partners under award numbers: U01DA041048, U01DA050989, U01DA051016, U01DA041022, U01DA051018, U01DA051037, U01DA050987, U01DA041174, U01DA041106, U01DA041117, U01DA041028, U01DA041134, U01DA050988, U01DA051039, U01DA041156, U01DA041025, U01DA041120, U01DA051038, U01DA041148, U01DA041093, U01DA041089, U24DA041123, U24DA041147. A full list of supporters is available at https://abcdstudy.org/about/federal-partners/. ABCD Consortium investigators designed and implemented the study and/or provided data but did not necessarily participate in the analysis or writing of this report. This manuscript reflects the views of the authors and may not reflect the opinions or views of the NIH or ABCD Consortium investigators. The ABCD data repository grows and changes over time. The ABCD data used in this report came from The ABCD Study® Release 6.1 (https://doi.org/10.82525/ab4q-qr87)].

DOIs can be found at https://www.nbdc-datahub.org/abcd-release-6-1. The authors wish to thank the youth and families participating in the ABCD Study® and all staff involved in data collection and curation.

## Declaration of Generative AI and AI-assisted technologies in the writing process

The authors used AI (Anthropic Claude) to reduce word count and correct grammar originally drafted by the authors. After using this tool, the authors reviewed and edited the content as needed and take full responsibility for the content of the publication.

## Declaration of Interests

Dr. Anders M. Dale is a Founding Director and holds equity in CorTechs Labs, Inc. (DBA Cortechs.ai), Precision Pro, Inc., Precision Health AS, Precision Health and Wellness, Inc., and Diploid Genomics, Inc. Dr. Dale is the President and a Board of Trustees member of the J. Craig Venter Institute (JCVI) and holds an appointment as Professor II at the University of Oslo in Norway. Dr. Kyung E. Rhee consults for Luro Health and receives stock options. No salary or direct research funding was received. The remaining authors have no conflicts of interest.

## Data Availability

ABCD Study® data release 6.1 is available for approved researchers in the NIH Brain Development Cohorts (NBDC) Data Hub (DOI: https://doi.org/10.82525/ab4q-qr87).

## Code Availability

The code to analyze the data and generate all figures of this manuscript is available on GitHub: https://github.com/Ali-Rigby/ABCD_bmi_microstructure

## Author Contributions

### Based on CRediT roles

Conceptualization; A.R., D.P., C.M., T.L.J, A.M.D

Data curation: A.R., D.P., J.L., R.L., D.J.H., K.E.R.

Formal analysis; A.R.

Funding acquisition; T.L.J., A.M.D.

Investigation: A.R., D.P., D.J.H., K.E.R.

Methodology; A.R., D.P., P.P., D.M.S., A.M.D.

Project administration; K.E.R., T.L.J., A.M.D.

Resources; J.L., A.M.D.

Software; A.R., D.P., P.P., D.M.S., A.M.D.

Supervision; D.P., P.P., C.M., T.L.J., A.M.D.

Visualization; A.R., D.P., P.P.

Roles/Writing -original draft; A.R., D.P.

Writing -review & editing. All authors

## Notes

### Summary of Updates

This version of the manuscript includes updated main text and supplementary revisions.

