## Supplement for "Assessing nonlinear associations between body mass index and brain microstructure across adolescence"

### Supplemental Material

**eMethods**

**eFigure 1. Distributions of Anthropometric Measures by Sex**

**eFigure 2. Distributions of Anthropometric Measures by ABCD Study Visit and Sex**

**eFigure 3. Distribution of Age**

**eFigure 4. Distribution of Puberty**

**eTable 1: Demographic Data for Matched Observations, Stratified by Scanner Software Group**

**eFigure 5. Distributions of Anthropometric Measures by Scanner Software Group**

**eFigure 6. Distributions of Anthropometric Measures by ABCD Study Visit and Scanner Software Group**

**eFigure 7. Distribution of Age by Scanner Software Group**

**eFigure 8. Distribution of Puberty by Scanner Software Group**

**eResults**

**eVideo 1: RNI Rate of Change Across the BMI Percentile Range (axial, females, X=69)**

**eVideo 2: RNI Rate of Change Across the BMI Percentile Range (axial, males, X=69)**

**eVideo 3: RNI Rate of Change Across the BMI Percentile Range (axial, females, X=91)**

**eVideo 4: RNI Rate of Change Across the BMI Percentile Range (axial, males, X=91)**

**eVideo 5: RNI Rate of Change Across the BMI Percentile Range (axial, females, X=97)**

**eVideo 6: RNI Rate of Change Across the BMI Percentile Range (axial, males, X=97)**

**eVideo 7: RNI Rate of Change Across the BMI Percentile Range (coronal, females, X=97)**

**eVideo 8: RNI Rate of Change Across the BMI Percentile Range (coronal, males, X=97)**

**eTable 2: Female-male Spatial Overlap of Voxelwise Associations**

**eFigure 9: Percent from Median BMI Voxelwise Maps**

**eVideo 9: RNI Rate of Change Across the Percent from Median BMI Range (axial, females, X=69)**

**eVideo 10: RNI Rate of Change Across the Percent from Median BMI Range (axial, males, X=69)**

**eVideo 11: RNI Rate of Change Across the Percent from Median BMI Range (axial, females, X=91)**

**eVideo 12: RNI Rate of Change Across the Percent from Median BMI Range (axial, males, X=91)**

**eVideo 13: RNI Rate of Change Across the Percent from Median BMI Range (axial, females, X=97)**

**eVideo 14: RNI Rate of Change Across the Percent from Median BMI Range (axial, males, X=97)**

**eVideo 15: RNI Rate of Change Across the Percent from Median BMI Range (coronal, females, X=97)**

**eVideo 16: RNI Rate of Change Across the Percent from Median BMI Range (coronal, males, X=97)**

**eFigure 10: BMI Percentile Voxelwise Maps by Scanner Software Group**

**eFigure 11: Percent from Median BMI Voxelwise Maps by Scanner Software Group**

**eFigure 12: Scanner-software-group-stratified BMI-RNI Associations in Selected Regions of Interest**

**eFigure 13: Scatterplots of BMI Percentile Beta Coefficient Estimates Between Scanner Software Group**

**eFigure 14: Scatterplots of Percent from Median Beta Coefficient Estimates Between Scanner Software Group**

**eReferences**

### eMethods:

##### ***Participants***

We analyzed data from the Adolescent Brain Cognitive Development℠ (ABCD) Study and included imaging visits from baseline and the 2-, 4-, and 6-year follow-up visits. The analytic sample comprised 21,629 observations (10,373 female; 11,256 male). Demographic characteristics of the analytic sample are summarized in Table 1.

##### ***Pubertal Status***

We assessed pubertal development using sex-appropriate Pubertal Development Scale (PDS)[1,2] category scores from both youth self-report and caregiver report. To address potential overestimation of pubertal status by youth and the caregiver's lack of awareness of a youth's status, we calculated a composite pubertal stage score as the mean of youth- and caregiver-reported PDS category scores, using whichever reporter was available when either was missing[3].

##### ***Covariate inclusion criteria***

We excluded visits if the absolute discrepancy between anthropometric assessment age and imaging visit age exceeded 100 days. Additional anthropometric QC removed visits with missing height/weight, biologically implausible decreasing height trajectories between visits, and extreme height, weight, and BMI values (using sex- and age-bin–specific 0.5th and 99.5th percentiles, with age bins of 8 to <10, 10 to <11, 11 to <12, 12 to <13, 13 to <14, 14 to <15, 15 to <16, and 16 to <18 years). For puberty scores, we checked for longitudinal consistency in the average composite score by examining the PDS score across consecutive imaging session pairs for regression (i.e., a decrease in the averaged PDS score at a later timepoint). For any such observed pair of values, we removed both observations. Imaging inclusion required the ABCD dMRI include indicator (mr_y_qc__incl__dmri_indicator) from the mr_y_qc__incl table. Design-matrix construction further required complete data on BMI phenotype, age, PDS, household income, caregiver education, genetic principal components (PCs) 1-10, and scanner/software. Distributions of BMI phenotypes, age, and puberty of our sample are visualized in eFigures 1-4.

**eFigure1.** Distributions of Anthropometric Measures by Sex

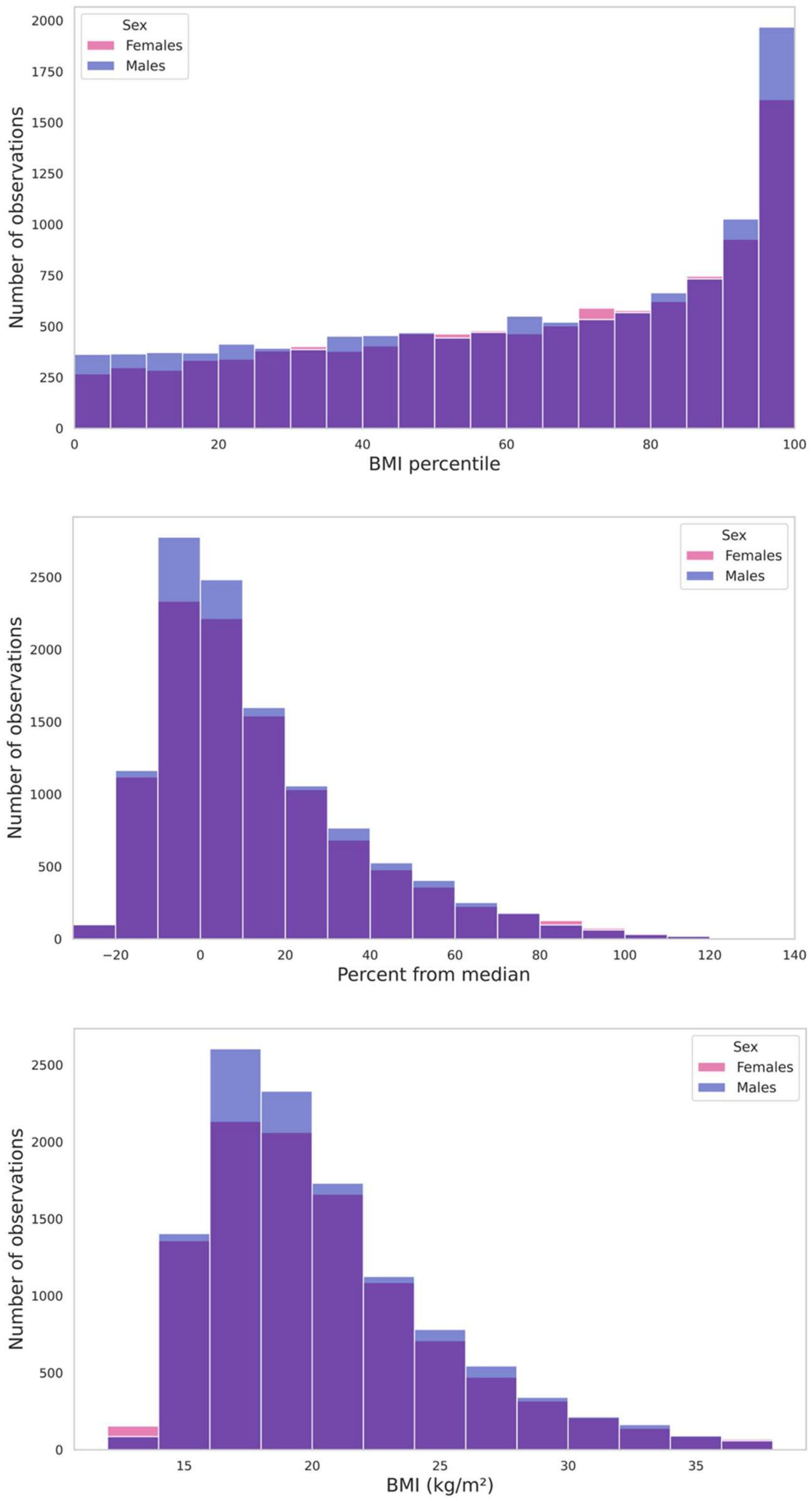

**eFigure 2.** Distributions of Anthropometric Measures by ABCD Study Visit and Sex

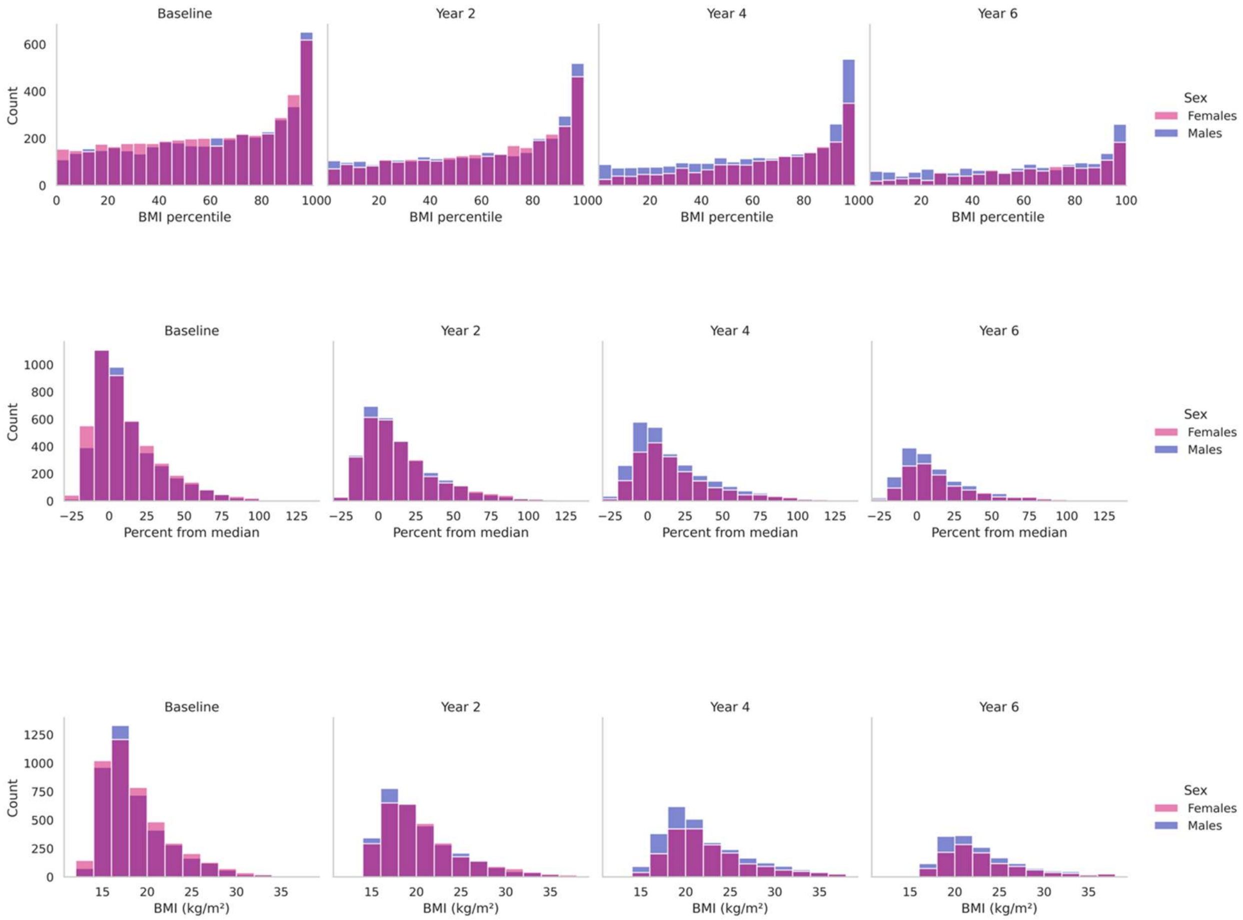

**eFigure 3.** Distribution of Age

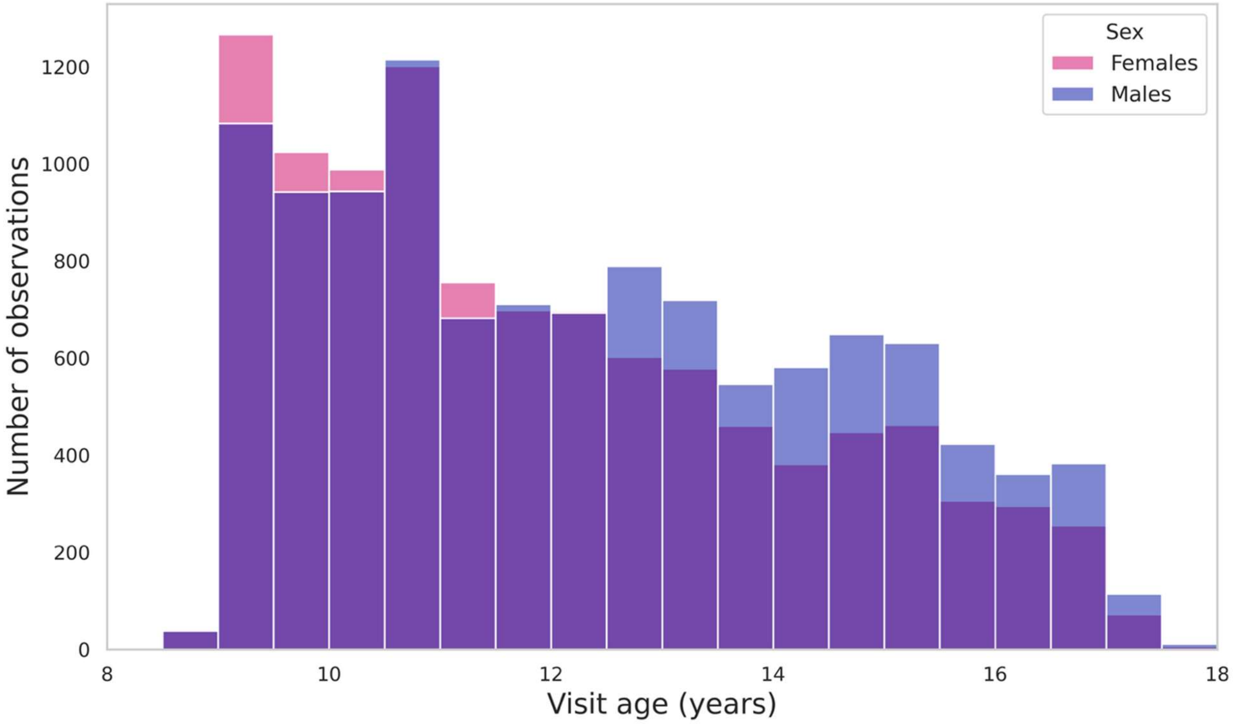


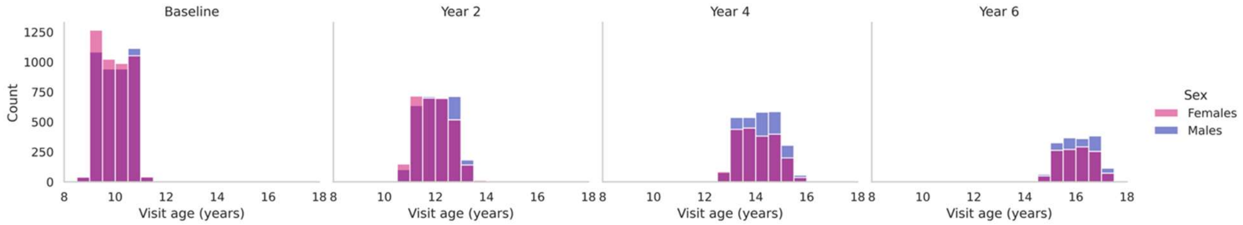

**eFigure 4.** Distribution of Puberty

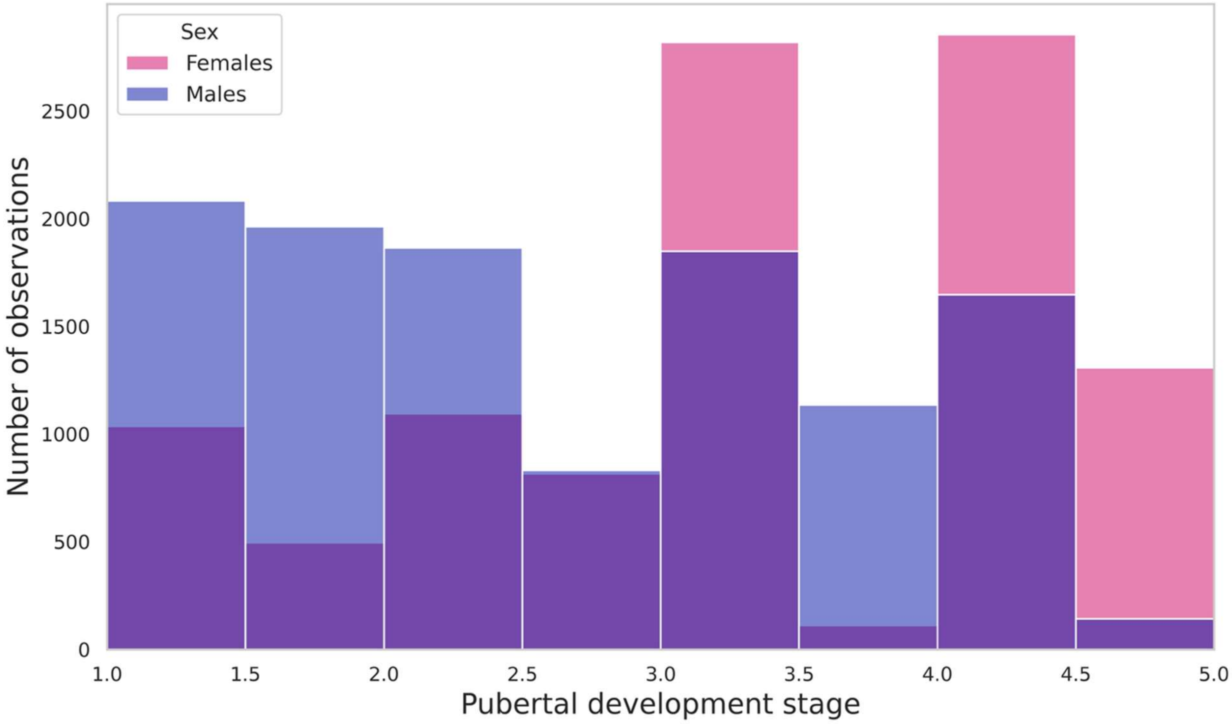


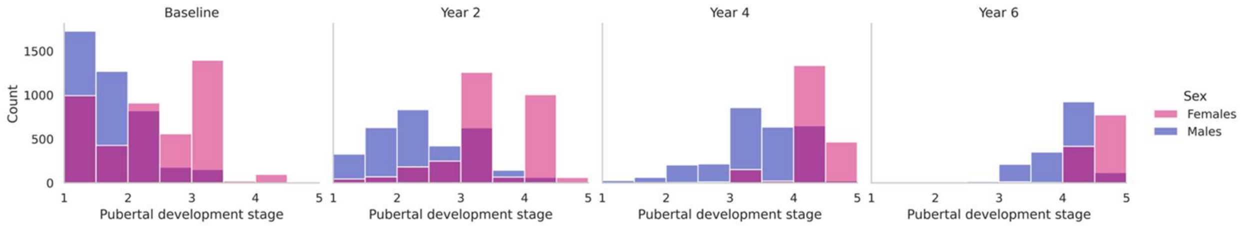

##### *Statistical models*

We estimated voxelwise associations with generalized additive mixed effects models using the Fast and Efficient Mixed-effects Analysis (FEMA) v4.0.3[4,5], fit separately in females and males. The conceptual model was:

$$\text{RNI} \sim 1 + s(\text{BMI phenotype}) + s(\text{age}) + s(\text{PDS}) + \text{PCs}_{1-10} + \text{income} + \text{education} + \text{scanner\_software} + (1|\text{family/subject}) + \varepsilon$$

where $s$ denotes a smooth function and $(1|\text{family/subject})$ denotes random intercepts for repeated measurements of subjects nested within families. We constructed nonlinear terms using natural cubic splines with unit heights at knots via singular value decomposition basis expansion without demeaning of design covariates[6,7]. We placed BMI phenotype knots at the sample 25th, 50th, and 75th percentiles of the BMI percentiles and percent from median BMI data (with boundary knots at the BMI range extremes) to concentrate flexibility where most observations fall. We placed age knots at the median values per visit, reflecting the longitudinal sampling structure. We placed puberty knots at integer category values 1-5 to correspond to the 5 stages of the PDS. Categorical covariates included household income, caregiver education, and scanner serial number + software version (reference levels were the mode). We accounted for population stratification by including the first 10 genetic principal components to account for population stratification.

##### *Voxelwise inference*

To test the association between BMI phenotype and RNI at each voxel, we performed an omnibus Wald test on the estimated spline basis coefficients, evaluating the joint null that all basis coefficients for the BMI smooth were simultaneously zero (no association between BMI and RNI) against the alternative that at least one was nonzero (a linear or nonlinear association). We controlled for multiple comparisons across brain voxels using the Benjamini-Hochberg false discovery rate (FDR) procedure at $q = 0.05$ (positive dependence setting)[8]. For visualization of maps, we used FDR correction to denote significance masks; we displayed uncorrected $-\log_{10}(p)$ values within FDR-significant voxels.

##### *Reconstruction of the BMI-RNI smooth*

At each voxel, the partial contribution of the BMI smooth ($\boldsymbol{B}$) to RNI was reconstructed as

$$\hat{f}(x) = \boldsymbol{B}(x)^{\top}\hat{\boldsymbol{\beta}}_{\text{spline}} + \hat{\beta}_0,$$

where $\boldsymbol{B}(x)$ are the BMI phenotype basis functions evaluated at $x$. This quantity is the partial effect of BMI on RNI (spline contribution plus intercept). We obtained voxelwise standard errors for $\hat{f}(x)$ from the corresponding coefficient covariance and the local rate of change of the association as the first derivative $d\hat{f}/dx$.

##### *Spatial Similarity of Sex-Stratified Maps*

To assess the similarity of the BMI-RNI association between sexes, we compared the male and female voxelwise maps. For each BMI phenotype, we quantified similarity in two ways: (1) the Pearson correlation, across voxels, between the male and female unthresholded omnibus Wald statistics from the test of the BMI smooth, indexing similarity of the continuous spatial pattern; and (2) the Dice similarity coefficient between the sets of voxels surviving FDR correction in each sex, defined as $2|F \cap M| / (|F| + |M|)$, indexing overlap of significant regions. These are descriptive measures of spatial similarity and do not constitute a formal test of sex differences.

##### *Region-of-interest analyses*

We extracted voxels corresponding to the top 9 most significant regions of interest (ROI) (based on the rank of the mean Wald statistic within each ROI). We used the FreeSurfer automated subcortical segmentation (aseg)[9] to define subcortical structures (nucleus accumbens, caudate, pallidum, putamen, and thalamus) and the AtlasTrack[10] probabilistic atlas to define white matter fiber tracts (anterior thalamic radiations, cingulate cingulum, and parahippocampal cingulum). We assigned voxels to an ROI if atlas probability exceeded 0.7 defined from the MINT probabilistic atlas[11], in which each voxel's value is the proportion of participants whose segmentation placed it in a given structure. We retained voxels assigned to the structure in more than 70% of participants (probability > 0.7), restricting analyses to confidently labeled voxels. We combined homologous left and right structures into bilateral ROIs where applicable and analyzed non-lateralized tracts such as forceps minor as single ROIs. Within each ROI, we averaged $\hat{f}(x)$ and $d\hat{f}/dx$ across voxels that were both inside the ROI mask and FDR-significant for the BMI-RNI Wald test. We formed confidence bands for ROI curves by averaging voxelwise 95% intervals for $\hat{f}(x)$.

##### *Gradients by CDC weight class*

To summarize how the BMI-RNI association changed across the BMI continuum, we averaged the ROI-averaged derivative curve over BMI-grid points falling within each CDC weight-class window, yielding a mean signed gradient (change in estimated RNI per BMI unit). Positive values indicate that the BMI-associated contribution to RNI increases with BMI within that range; negative values indicate a decrease. For BMI percentile models, the CDC weight-class windows were set as the fixed percentile bands defining underweight, healthy weight, overweight, and obesity. For percent from median BMI models, we mapped the weight class windows using the minimum and maximum observed percent from median values corresponding to the BMI percentile definitions. Gradient values in tables are reported $\times 10^{-5}$.

##### *Sensitivity Analysis*

We repeated our voxelwise analyses to assess the effect of software version on the BMI-RNI associations. We stratified our analyses into groups of data acquired on VE11 software and all other scanner-software versions (non-VE11). Since VE11 visits outnumbered non-VE11 visits, we matched based on BMI and age (see eTable1 and eFigures 5-8). The matched sample comprised 7,889 VE11 visits (3,725 females, 4,164 males) and 7,579 non-VE11 visits (3,727 females, 3, 852 males).

We recomputed spline basis functions for BMI phenotype, age, and puberty on the combined VE11 and non-VE11 sample. We fit generalized additive mixed effects models for each scanner software group using the same model as our primary analyses. The only difference between the two models involved covarying for scanner serial number and software version in the non-VE11 models, while the VE11 model only corrected for scanner serial number, since all scans shared the same software version. Omnibus tests and ROI analyses followed the same procedures as described above in primary analyses.

| **Characteristic**[a] | **Overall** (n = 15,468 observations) | **VE11** (n = 7,889 observations) | **noVE11** (n = 7,579 observations) | **p-value** |
|---|---|---|---|---|
| **Sex, No. (%)** | | | | 0.016 |
| Female | 7,452 (48.2%) | 3,725 (47.2%) | 3,727 (49.2%) | |
| Male | 8,016 (51.8%) | 4,164 (52.8%) | 3,852 (50.8%) | |
| **Age, y** | 12.2 (2.3) | 12.3 (2.3) | 12.2 (2.3) | 0.198 |
| **BMI, kg/m^2** | 20.8 (4.8) | 20.8 (4.8) | 20.7 (4.8) | 0.369 |
| **CDC extended BMI percentile**[b] | 63.7 (29.1) | 63.9 (29.1) | 63.6 (29.1) | 0.565 |
| **Percent from median BMI, age/sex** | 14.5 (24.3) | 14.6 (24.4) | 14.3 (24.2) | 0.583 |
| **Puberty** | 2.8 (1.1) | 2.8 (1.1) | 2.7 (1.1) | 0.161 |
| **CDC weight class, No. (%)**[d] | | | | 0.911 |
| Underweight | 332 (2.1%) | 170 (2.2%) | 162 (2.1%) | |
| Healthy weight | 9,949 (64.3%) | 5,056 (64.1%) | 4,893 (64.6%) | |
| Overweight | 2,587 (16.7%) | 1,321 (16.7%) | 1,266 (16.7%) | |
| Obesity | 2,600 (16.8%) | 1,342 (17.0%) | 1,258 (16.6%) | |
| **Study wave, No. (%)** | | | | 0.007 |
| Baseline | 5,812 (37.6%) | 2,994 (38.0%) | 2,818 (37.2%) | |
| 2-Year FU | 4,142 (26.8%) | 2,079 (26.4%) | 2,063 (27.2%) | |
| 4-Year FU | 3,296 (21.3%) | 1,742 (22.1%) | 1,554 (20.5%) | |
| 6-Year FU | 2,218 (14.3%) | 1,074 (13.6%) | 1,144 (15.1%) | |
| **Race, No. (%)**[c] | | | | <0.001 |
| White | 10,724 (69.3%) | 5,661 (71.8%) | 5,063 (66.8%) | |
| Black or African American | 1,934 (12.5%) | 1,237 (15.7%) | 697 (9.2%) | |
| Asian | 341 (2.2%) | 96 (1.2%) | 245 (3.2%) | |
| American Indian or Alaska Native | 74 (0.5%) | 20 (0.3%) | 54 (0.7%) | |
| Native Hawaiian or Other Pacific Islander | 23 (0.1%) | 16 (0.2%) | 7 (0.1%) | |
| More than 1 race | 1,758 (11.4%) | 690 (8.7%) | 1,068 (14.1%) | |
| Other or unknown | 614 (4.0%) | 169 (2.1%) | 445 (5.9%) | |
| **Ethnicity, No. (%)**[d] | | | | <0.001 |
| Hispanic | 3,127 (20.2%) | 1,363 (17.3%) | 1,764 (23.3%) | |
| Non-Hispanic | 12,299 (79.5%) | 6,503 (82.4%) | 5,796 (76.5%) | |
| Missing | 42 (0.3%) | 23 (0.3%) | 19 (0.3%) | |

| | | | | |
|---|---|---|---|---|
| **Annual household income, No. (%)** | | | | <0.001 |
| <$25 000 | 1,501 (9.7%) | 724 (9.2%) | 777 (10.3%) | |
| $25 000-$49 999 | 1,801 (11.6%) | 829 (10.5%) | 972 (12.8%) | |
| $50 000-$74 999 | 1,811 (11.7%) | 975 (12.4%) | 836 (11.0%) | |
| $75 000-$99 999 | 2,005 (13.0%) | 1,094 (13.9%) | 911 (12.0%) | |
| $100 000-$199 999 | 4,975 (32.2%) | 2,611 (33.1%) | 2,364 (31.2%) | |
| ≥$200 000 | 2,227 (14.4%) | 1,067 (13.5%) | 1,160 (15.3%) | |
| Declined to answer | 601 (3.9%) | 323 (4.1%) | 278 (3.7%) | |
| Don't know | 547 (3.5%) | 266 (3.4%) | 281 (3.7%) | |
| **Highest parental education, No. (%)** | | | | <0.001 |
| Up to high school (no diploma) | 687 (4.4%) | 234 (3.0%) | 453 (6.0%) | |
| High school diploma/GED | 1,246 (8.1%) | 637 (8.1%) | 609 (8.0%) | |
| Some college | 4,358 (28.2%) | 2,341 (29.7%) | 2,017 (26.6%) | |
| Bachelor's degree | 3,587 (23.2%) | 1,952 (24.7%) | 1,635 (21.6%) | |
| Graduate or professional degree | 5,590 (36.1%) | 2,725 (34.5%) | 2,865 (37.8%) | |

**eTable 1: Demographic Data for Matched Observations, Stratified by Scanner Software Group**

[a] Categorical values are presented as No. (%); continuous variables are presented as mean (SD).

[b] CDC extended BMI percentile was derived using CDC age- and sex-specific growth reference data. CDC weight classes were derived from CDC extended BMI percentile as underweight (<5th percentile), healthy weight (5th to <85th percentile), overweight (85th to <95th percentile), and obesity (>=95th percentile).

[c] Race (NIH 7 categories) was self-reported by the caregiver. Descriptive statistics are reported for self-reported race only for interpretation of sample diversity.

[d] Ethnicity (2 categories) was self-reported by the caregiver.

[e] P values compare VE11 vs noVE11 at the visit level using Wilcoxon rank-sum tests (Mann-Whitney U) for continuous variables and chi-squared tests for categorical variables.

Abbreviations: BMI, body mass index; CDC, Centers for Disease Control and Prevention; GED, Generalized Education Degree; SD, standard deviation.

**eFigure 5.** Distributions of Anthropometric Measures by Scanner Software Group

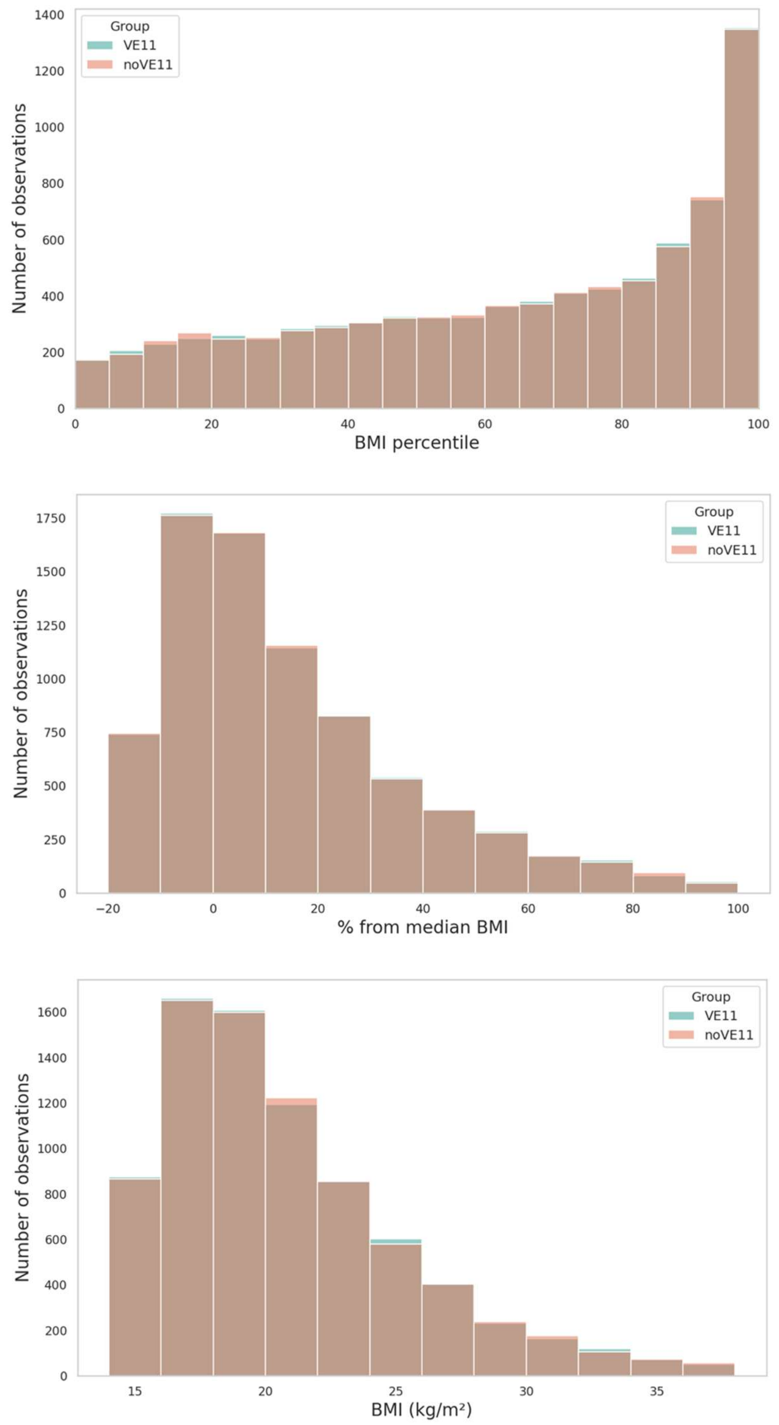

**eFigure 6.** Distributions of Anthropometric Measures by ABCD Study Visit and Scanner Software Group

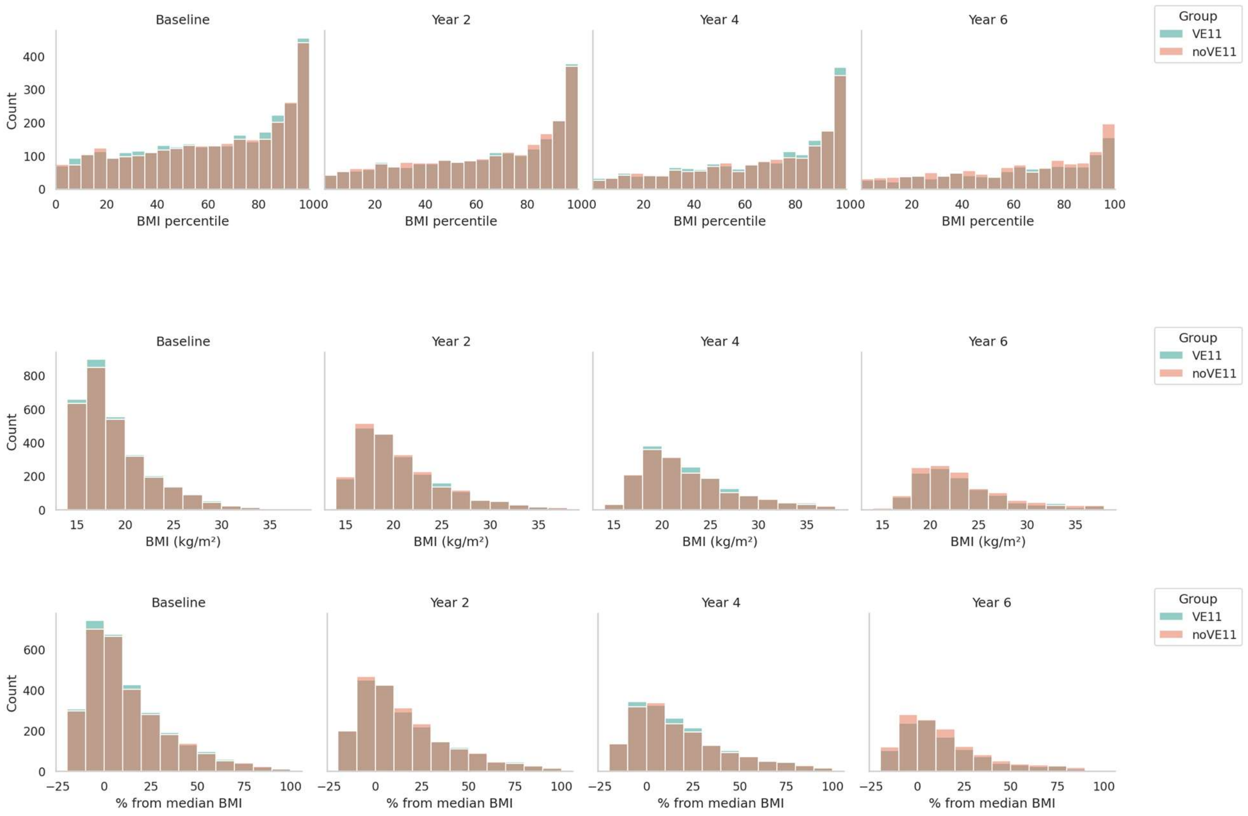

**eFigure 7.** Distribution of Age by Scanner Software Group

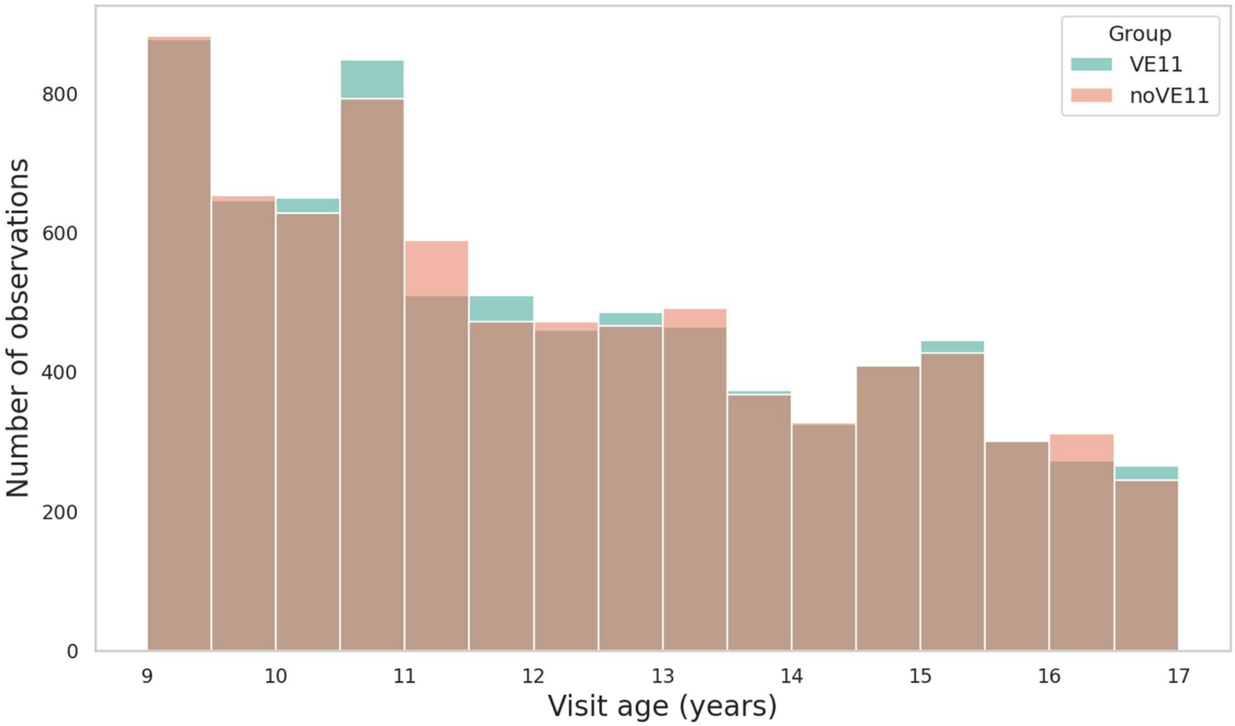


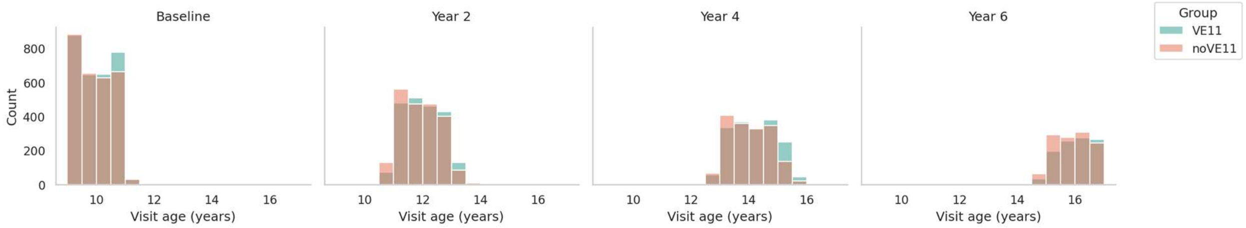

**eFigure 8.** Distribution of Puberty by Scanner Software Group

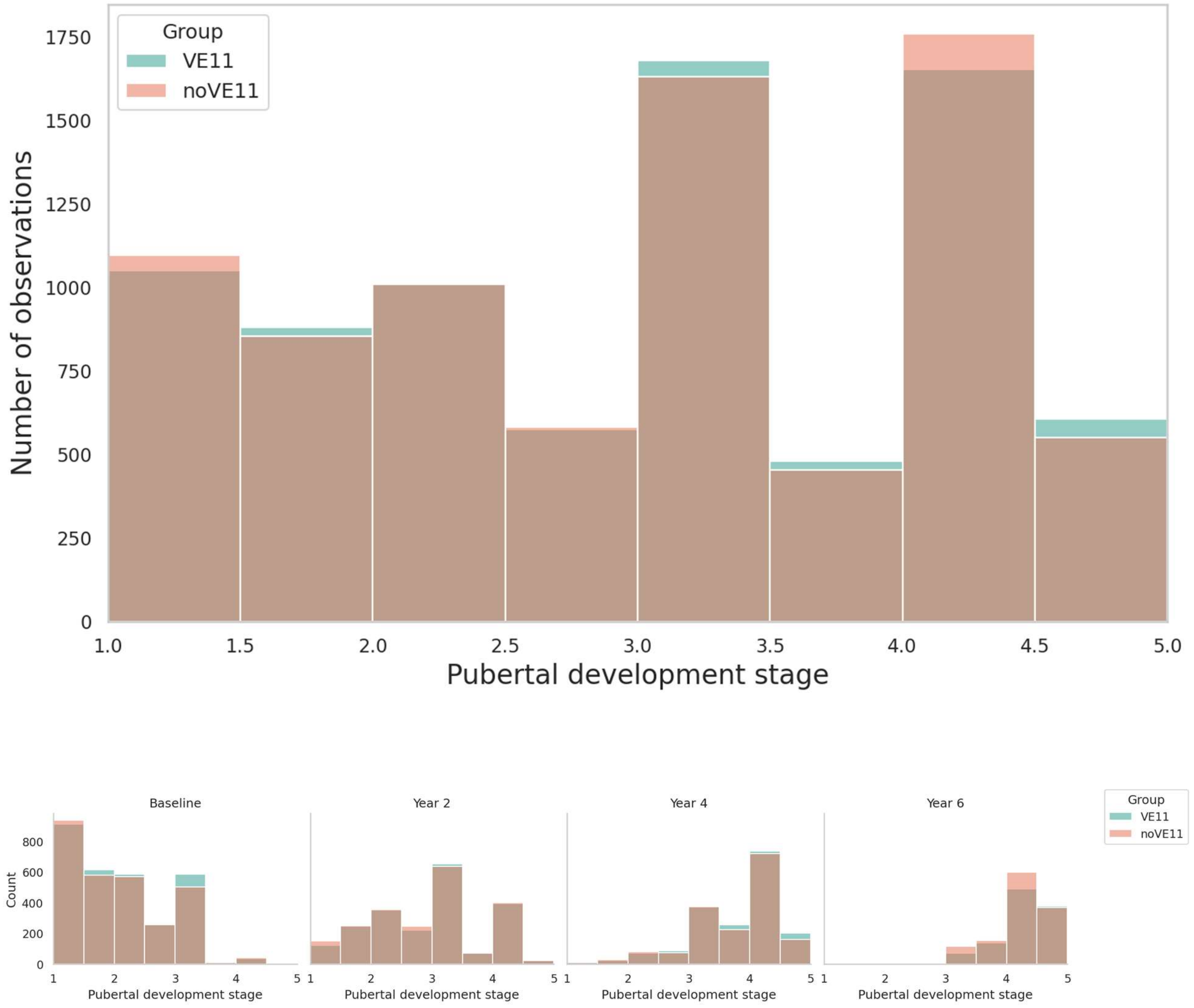


#### ***Software***

Data preprocessing, analyses and visualizations were implemented in R version 4.4.3[12], MATLAB R2022a[13], and Python 3.13.13[14]. R packages included arrow (24.0.0), dplyr (1.2.1), tidyverse (2.0.0), data.table (1.17.8), plyr (1.8.9), gtsummary (2.5.1), and gt (1.3.0). Python packages included NumPy (2.3.3), pandas (2.3.3), matplotlib (3.10.7), seaborn (0.13.2), SciPy (1.16.2), and h5py (3.15.0). The code for FEMA is publicly available at https://github.com/cmig-research-group/cmig_tools.

##### *Code Availability*

The code to analyze the data and generate all figures of this manuscript is available on GitHub: https://github.com/Ali-Rigby/ABCD_bmi_microstructure

### eResults:

**eVideo 1: RNI Rate of Change Across the BMI Percentile Range (axial, females, X=69)**

Axial voxelwise maps of the first derivative of RNI with respect to BMI percentile, across the full range, in females; slice at atlas coordinates (X=69, Y=100, Z=147), showing the cingulum and forceps minor; warmer colors indicate a greater rate of change; MP4 format.

**eVideo 2: RNI Rate of Change Across the BMI Percentile Range (axial, males, X=69)**

Axial voxelwise maps of the first derivative of RNI with respect to BMI percentile, across the full range, in males; slice at atlas coordinates (X=69, Y=100, Z=147), showing the cingulum and forceps minor; warmer colors indicate a greater rate of change; MP4 format.

**eVideo 3: RNI Rate of Change Across the BMI Percentile Range (axial, females, X=91)**

Axial voxelwise maps of the first derivative of RNI with respect to BMI percentile, across the full range, in females; slice at atlas coordinates (X=91, Y=100, Z=147), showing the anterior thalamic radiations, caudate, forceps minor, pallidum, putamen, and thalamus; warmer colors indicate a greater rate of change; MP4 format.

**eVideo 4: RNI Rate of Change Across the BMI Percentile Range (axial, males, X=91)**

Axial voxelwise maps of the first derivative of RNI with respect to BMI percentile, across the full range, in males; slice at atlas coordinates (X=91, Y=100, Z=147), showing the anterior thalamic radiations, caudate, forceps minor, pallidum, putamen, and thalamus; warmer colors indicate a greater rate of change; MP4 format.

**eVideo 5: RNI Rate of Change Across the BMI Percentile Range (axial, females, X=97)**

Axial voxelwise maps of the first derivative of RNI with respect to BMI percentile, across the full range, in females; slice at atlas coordinates (X=97, Y=100, Z=147), showing the accumbens, anterior thalamic radiations, forceps minor, pallidum, putamen, and thalamus; warmer colors indicate a greater rate of change; MP4 format.

**eVideo 6: RNI Rate of Change Across the BMI Percentile Range (axial, males, X=97)**

Axial voxelwise maps of the first derivative of RNI with respect to BMI percentile, across the full range, in males; slice at atlas coordinates (X=97, Y=100, Z=147), showing the accumbens, anterior thalamic radiations, forceps minor, pallidum, putamen, and thalamus; warmer colors indicate a greater rate of change; MP4 format.

**eVideo 7: RNI Rate of Change Across the BMI Percentile Range (coronal, females, X=97)**

Coronal voxelwise maps of the first derivative of RNI with respect to BMI percentile, across the full range, in females; slice at atlas coordinates (X=97, Y=100, Z=147), showing the nucleus accumbens, anterior thalamic radiations, caudate, and putamen; warmer colors indicate a greater rate of change; MP4 format.

**eVideo 8: RNI Rate of Change Across the BMI Percentile Range (coronal, males, X=97)**

Coronal voxelwise maps of the first derivative of RNI with respect to BMI percentile, across the full range, in males; slice at atlas coordinates (X=97, Y=100, Z=147), showing the nucleus accumbens, anterior thalamic radiations, caudate, and putamen; warmer colors indicate a greater rate of change; MP4 format.

| Phenotype | Pearson correlation | Dice coefficient (FDR) |
| --- | --- | --- |
| **BMI percentile** | 0.883 | 0.822 |
| **Percent from median** | 0.891 | 0.839 |

**eTable 2. Female-male spatial overlap of voxelwise associations**

**Note.** Pearson is for unsigned Wald $W_s$ and Dice is for FDR maps ($q = 0.05$).

**eFigure 9.** Percent from median BMI voxelwise maps

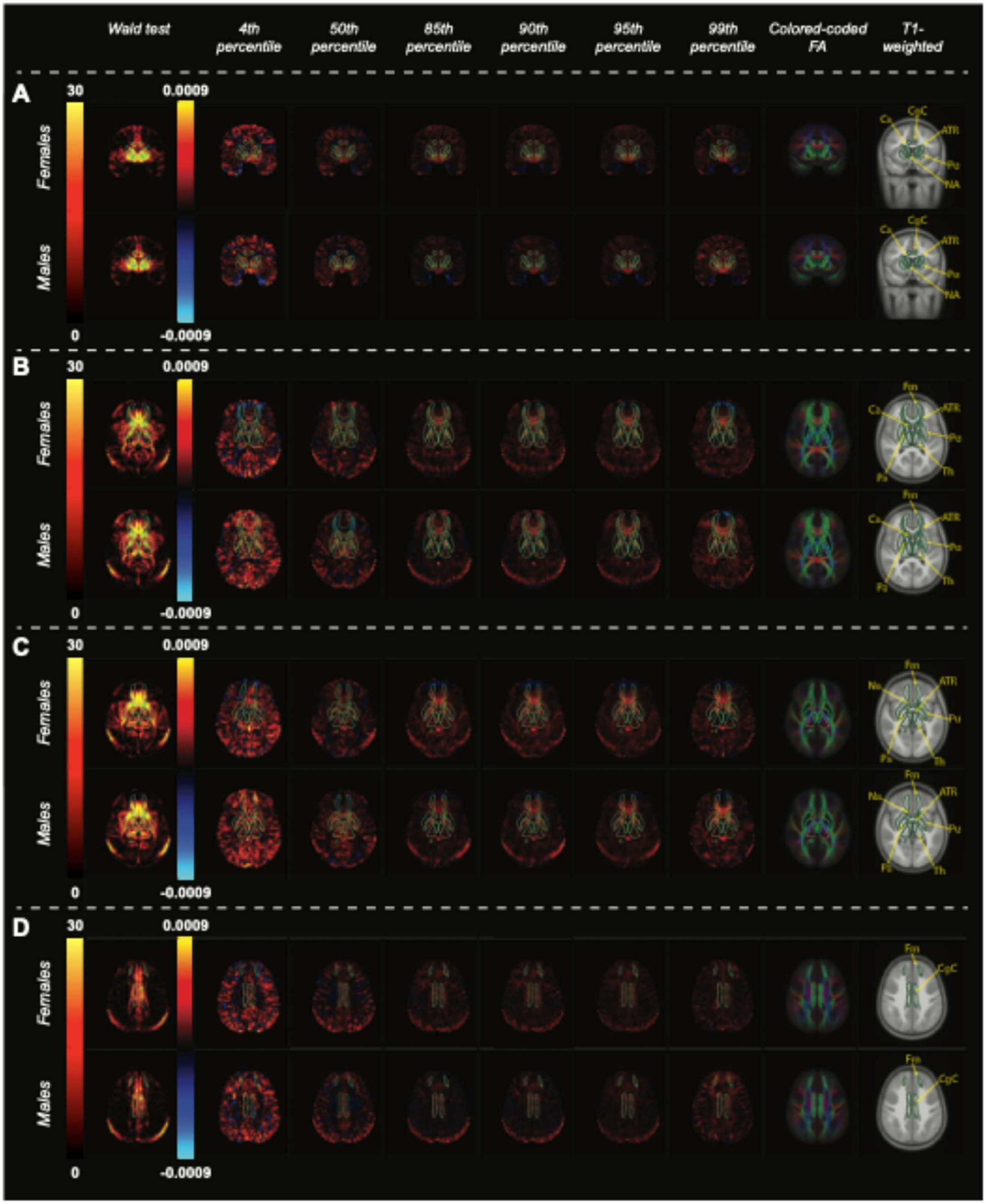

**Note. Voxelwise BMI-RNI associations and local rates of change across the percent from median BMI range.**

Column 1. Voxelwise $-\log_{10}$ p-value maps from the omnibus Wald test of the nonlinear percent from median BMI effect on restricted normalized isotropic signal fraction (RNI), thresholded at FDR $q < 0.05$. Columns 2-7. Derivative maps of the BMI-RNI smooth, showing the estimated instantaneous rate of change in the BMI effect on RNI per unit percent from median BMI corresponding to BMI percentile values for the 4th (underweight), 50th (healthy weight), 85th & 90th (overweight), and 95th & 99th (obesity) BMI percentiles. (The columns are labeled as BMI percentile because percent from median BMI values differ between females and males.) Warm colors indicate that the BMI-associated contribution to RNI increases with percent from median BMI at that location; cool colors indicate a decrease. Columns 8-9. Color-coded fractional anisotropy (left-right: red, anterior-posterior: green, inferior-superior: blue) and T1-weighted images for anatomical reference. (A) Coronal view highlighting the nucleus accumbens, anterior thalamic radiations, caudate, and putamen. (B) Axial view highlighting the caudate, forceps minor, pallidum, putamen, and thalamus. (C) Axial view highlighting the accumbens, anterior thalamic radiations, forceps minor, pallidum, putamen, and thalamus. (D) Axial view highlighting the cingulate cingulum. Subcortical ROI outlines and labels are shown from the FreeSurfer automated subcortical segmentation and AtlasTrack. All maps were derived from sex-stratified voxelwise linear mixed-effects models including smooth terms for percent from median BMI, age, and pubertal development, with covariates for genetic principal components, household income, highest parental education, and scanner/software version. The bright band at the posterior of the brain on axial slices is a chemical-shift (fat) artifact and does not reflect brain microstructure. Label abbreviations: NA = nucleus accumbens, ATR = anterior thalamic radiations; Ca = caudate; CgC = cingulate cingulum; Fm = forceps minor; Pa = pallidum; Pu = putamen; Th = thalamus.

**eVideo 9: RNI Rate of Change Across the Percent from Median BMI Range (axial, females, X=69)**

Axial voxelwise maps of the first derivative of RNI with respect to percent from median BMI, across the full range, in females; slice at atlas coordinates (X=69, Y=100, Z=147), showing the cingulum and forceps minor; warmer colors indicate a greater rate of change; MP4 format.

**eVideo 10: RNI Rate of Change Across the Percent from Median BMI Range (axial, males, X=69)**

Axial voxelwise maps of the first derivative of RNI with respect to percent from median BMI, across the full range, in males; slice at atlas coordinates (X=69, Y=100, Z=147), showing the cingulum and forceps minor; warmer colors indicate a greater rate of change; MP4 format.

**eVideo 11: RNI Rate of Change Across the Percent from Median BMI Range (axial, females, X=91)**

Axial voxelwise maps of the first derivative of RNI with respect to percent from median BMI, across the full range, in females; slice at atlas coordinates (X=91, Y=100, Z=147), showing the anterior thalamic radiations, caudate, forceps minor, pallidum, putamen, and thalamus; warmer colors indicate a greater rate of change; MP4 format.

**eVideo 12: RNI Rate of Change Across the Percent from Median BMI Range (axial, males, X=91)**

Axial voxelwise maps of the first derivative of RNI with respect to percent from median BMI, across the full range, in males; slice at atlas coordinates (X=91, Y=100, Z=147), showing the anterior thalamic radiations, caudate, forceps minor, pallidum, putamen, and thalamus; warmer colors indicate a greater rate of change; MP4 format.

**eVideo 13: RNI Rate of Change Across the Percent from Median BMI Range (axial, females, X=97)**

Axial voxelwise maps of the first derivative of RNI with respect to percent from median BMI, across the full range, in females; slice at atlas coordinates (X=97, Y=100, Z=147), showing the accumbens, anterior thalamic radiations, forceps minor, pallidum, putamen, and thalamus; warmer colors indicate a greater rate of change; MP4 format.

**eVideo 14: RNI Rate of Change Across the Percent from Median BMI Range (axial, males, X=97)**

Axial voxelwise maps of the first derivative of RNI with respect to percent from median BMI, across the full range, in males; slice at atlas coordinates (X=97, Y=100, Z=147), showing the accumbens, anterior thalamic radiations, forceps minor, pallidum, putamen, and thalamus; warmer colors indicate a greater rate of change; MP4 format.

**eVideo 15: RNI Rate of Change Across the Percent from Median BMI Range (coronal, females, X=97)**

Coronal voxelwise maps of the first derivative of RNI with respect to percent from median BMI, across the full range, in females; slice at atlas coordinates (X=97, Y=100, Z=147), showing the nucleus accumbens, anterior thalamic radiations, caudate, and putamen; warmer colors indicate a greater rate of change; MP4 format.

**eVideo 16: RNI Rate of Change Across the Percent from Median BMI Range (coronal, males, X=97)**

Coronal voxelwise maps of the first derivative of RNI with respect to percent from median BMI, across the full range, in males; slice at atlas coordinates (X=97, Y=100, Z=147), showing the nucleus accumbens, anterior thalamic radiations, caudate, and putamen; warmer colors indicate a greater rate of change; MP4 format.

##### *Sensitivity Analysis by Scanner Software Group*

Both VE11 and non-VE11 groups showed similar spatial distribution and effect sizes of BMI phenotype-RNI associations compared with the main results (see eFigures 10-16), indicating that the associations are not driven by the potential confound. However, we observed subtle differences: Voxelwise maps revealed that the ATR, forceps minor, pallidum, and thalamus showed greater Wald statistics in the non-VE11 sample compared to the VE11 sample (eFigures 10 & 11). The opposite effect was observed in caudate, nucleus accumbens, and putamen (eFigures 10 & 11). For most ROIs, the BMI phenotype-RNI associations were slightly greater in the non-VE11 sample, with wider confidence intervals (eFigure 12). These differences may be attributable to greater scanner/software heterogeneity in the non-VE11 sample, sample size differences, and residual differences in sample composition. Overall, these results are in agreement with our primary analyses.

**eFigure 10: BMI Percentile Voxelwise Maps by Scanner Software Group**

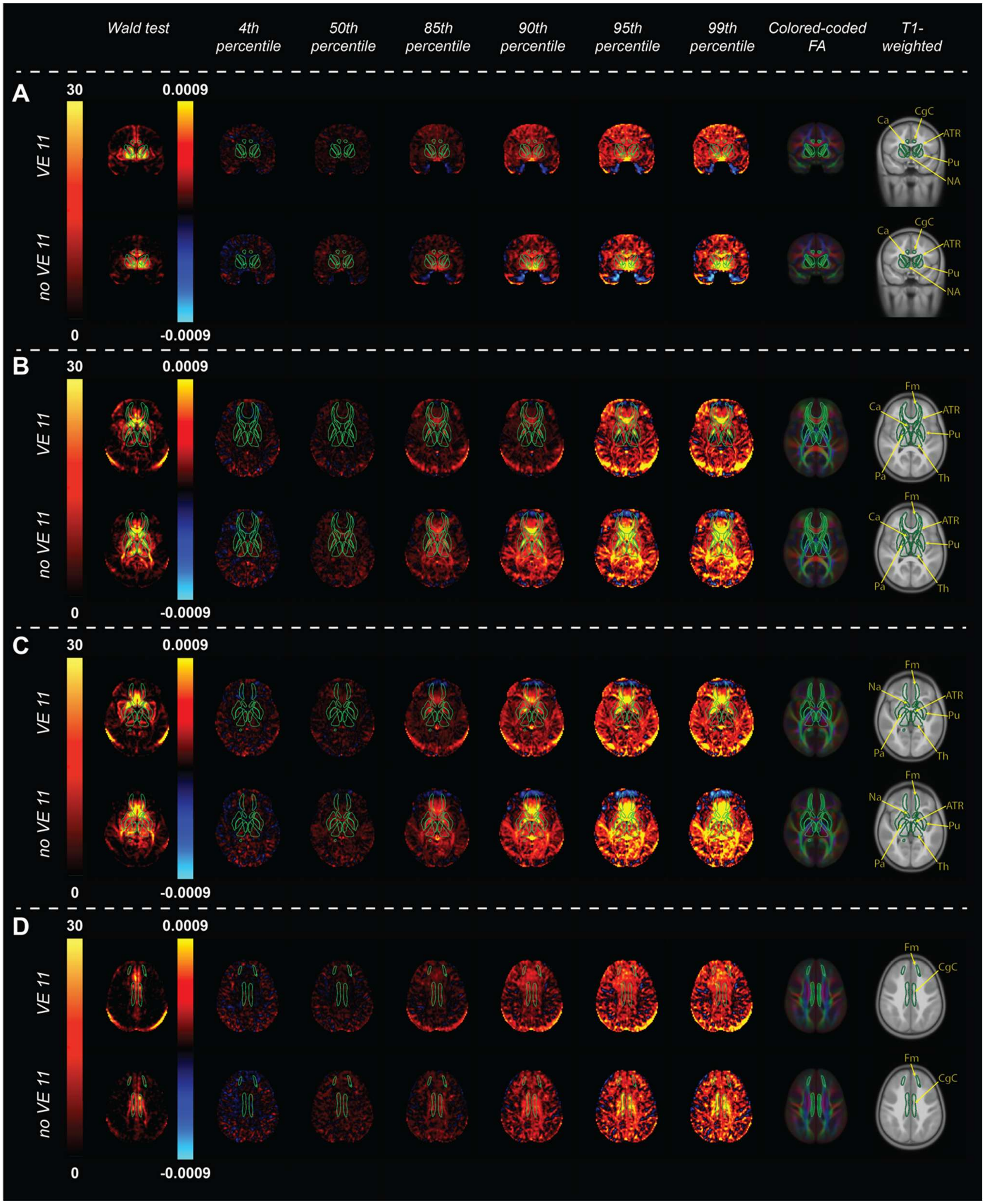

**Note. Voxelwise BMI-RNI associations and local rates of change across the BMI percentile range for VE11 and non-VE11 samples.**

Column 1. Voxelwise $-\log_{10}$ p-value maps from the omnibus Wald test of the nonlinear BMI percentile effect on restricted normalized isotropic signal fraction (RNI), thresholded at FDR $q < 0.05$. Columns 2-7. Derivative maps of the BMI-RNI smooth, showing the estimated instantaneous rate of change in the BMI effect on RNI per unit percent from median BMI corresponding to BMI percentile values for the 4th (underweight), 50th (healthy weight), 85th & 90th (overweight), and 95th & 99th (obesity) BMI percentiles. Warm colors indicate that the BMI-associated contribution to RNI increases with BMI percentile at that location; cool colors indicate a decrease. Columns 8-9. Color-coded fractional anisotropy (left-right: red, anterior-posterior: green, inferior-superior: blue) and T1-weighted images for anatomical reference. (A) Coronal view highlighting the nucleus accumbens, anterior thalamic radiations, caudate, and putamen. (B) Axial view highlighting the caudate, forceps minor, pallidum, putamen, and thalamus. (C) Axial view highlighting the accumbens, anterior thalamic radiations, forceps minor, pallidum, putamen, and thalamus. (D) Axial view highlighting the cingulate cingulum. Subcortical ROI outlines and labels are shown from the FreeSurfer automated subcortical segmentation and AtlasTrack. All maps were derived from scanner-software-group-stratified voxelwise linear mixed-effects models including smooth terms for BMI percentile, age, and pubertal development, with covariates for genetic principal components, household income, highest parental education, and scanner/software (non-VE11) or scanner (VE11) version. The bright band at the posterior of the brain on axial slices is a chemical-shift (fat) artifact and does not reflect brain microstructure. Label abbreviations: NA = nucleus accumbens, ATR = anterior thalamic radiations; Ca = caudate; CgC = cingulate cingulum; Fm = forceps minor; Pa = pallidum; Pu = putamen; Th = thalamus.

**eFigure 11: Percent from Median BMI Voxelwise Maps by Scanner Software Group**

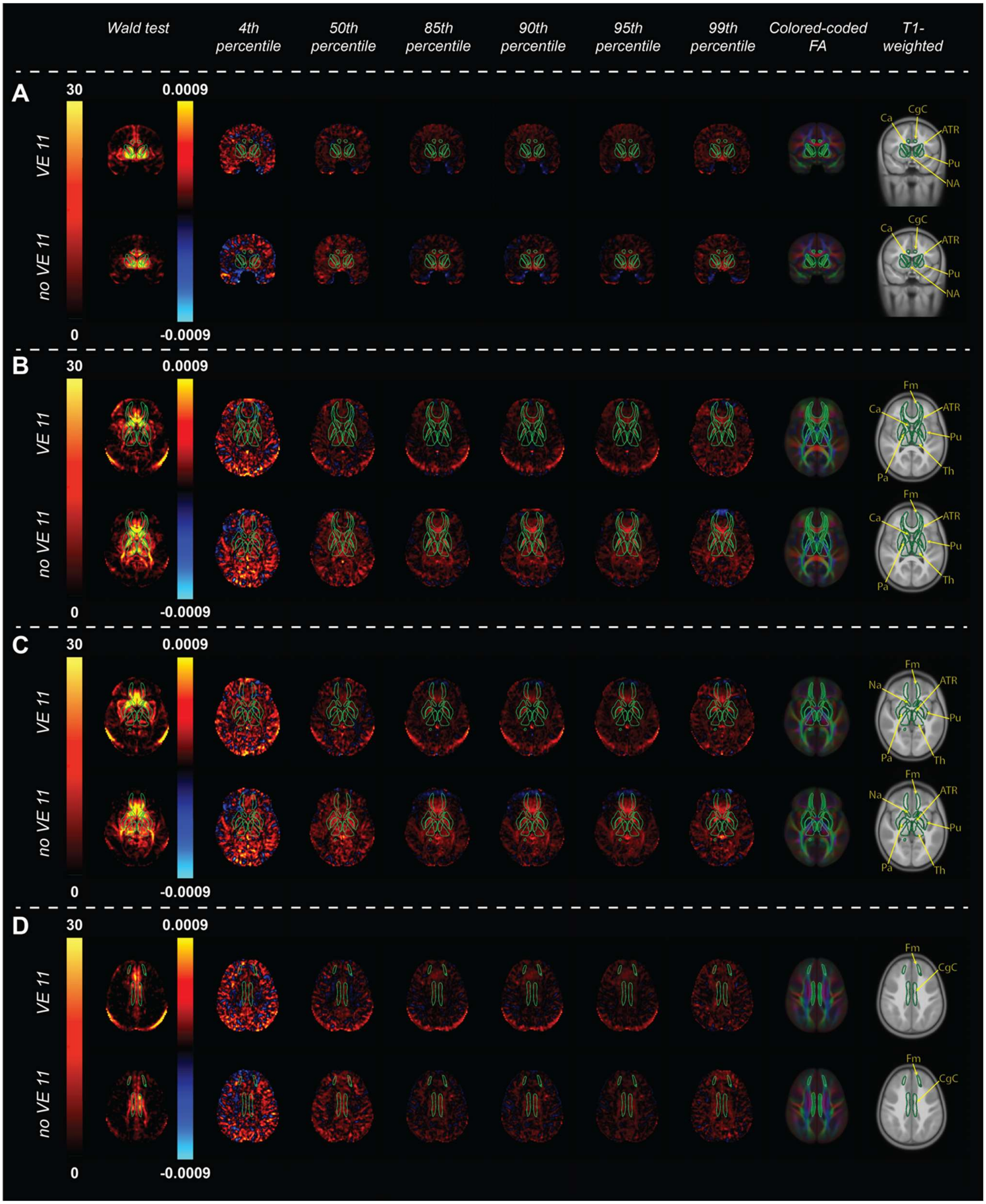

**Note. Voxelwise BMI-RNI associations and local rates of change across the percent from median BMI range for VE11 and non-VE11 samples.**

Column 1. Voxelwise $-\log_{10}$ p-value maps from the omnibus Wald test of the nonlinear percent from median BMI effect on restricted normalized isotropic signal fraction (RNI), thresholded at FDR $q < 0.05$. Columns 2-7. Derivative maps of the BMI-RNI smooth, showing the estimated instantaneous rate of change in the BMI effect on RNI per unit percent from median BMI corresponding to BMI percentile values for the 4th (underweight), 50th (healthy weight), 85th & 90th (overweight), and 95th & 99th (obesity) BMI percentiles. Warm colors indicate that the BMI-associated contribution to RNI increases with percent from median BMI at that location; cool colors indicate a decrease. Columns 8-9. Color-coded fractional anisotropy (left-right: red, anterior-posterior: green, inferior-superior: blue) and T1-weighted images for anatomical reference. (A) Coronal view highlighting the nucleus accumbens, anterior thalamic radiations, caudate, and putamen. (B) Axial view highlighting the caudate, forceps minor, pallidum, putamen, and thalamus. (C) Axial view highlighting the accumbens, anterior thalamic radiations, forceps minor, pallidum, putamen, and thalamus. (D) Axial view highlighting the cingulate cingulum. Subcortical ROI outlines and labels are shown from the FreeSurfer automated subcortical segmentation and AtlasTrack. All maps were derived from scanner-software-group-stratified voxelwise linear mixed-effects models including smooth terms for percent from median BMI, age, and pubertal development, with covariates for genetic principal components, household income, highest parental education, and scanner/software (non-VE11) or scanner (VE11) version. The bright band at the posterior of the brain on axial slices is a chemical-shift (fat) artifact and does not reflect brain microstructure. Label abbreviations: NA = nucleus accumbens, ATR = anterior thalamic radiations; Ca = caudate; CgC = cingulate cingulum; Fm = forceps minor; Pa = pallidum; Pu = putamen; Th = thalamus.

**eFigure 12: Scanner-software-group-stratified BMI-RNI Associations in Selected Regions of Interest**

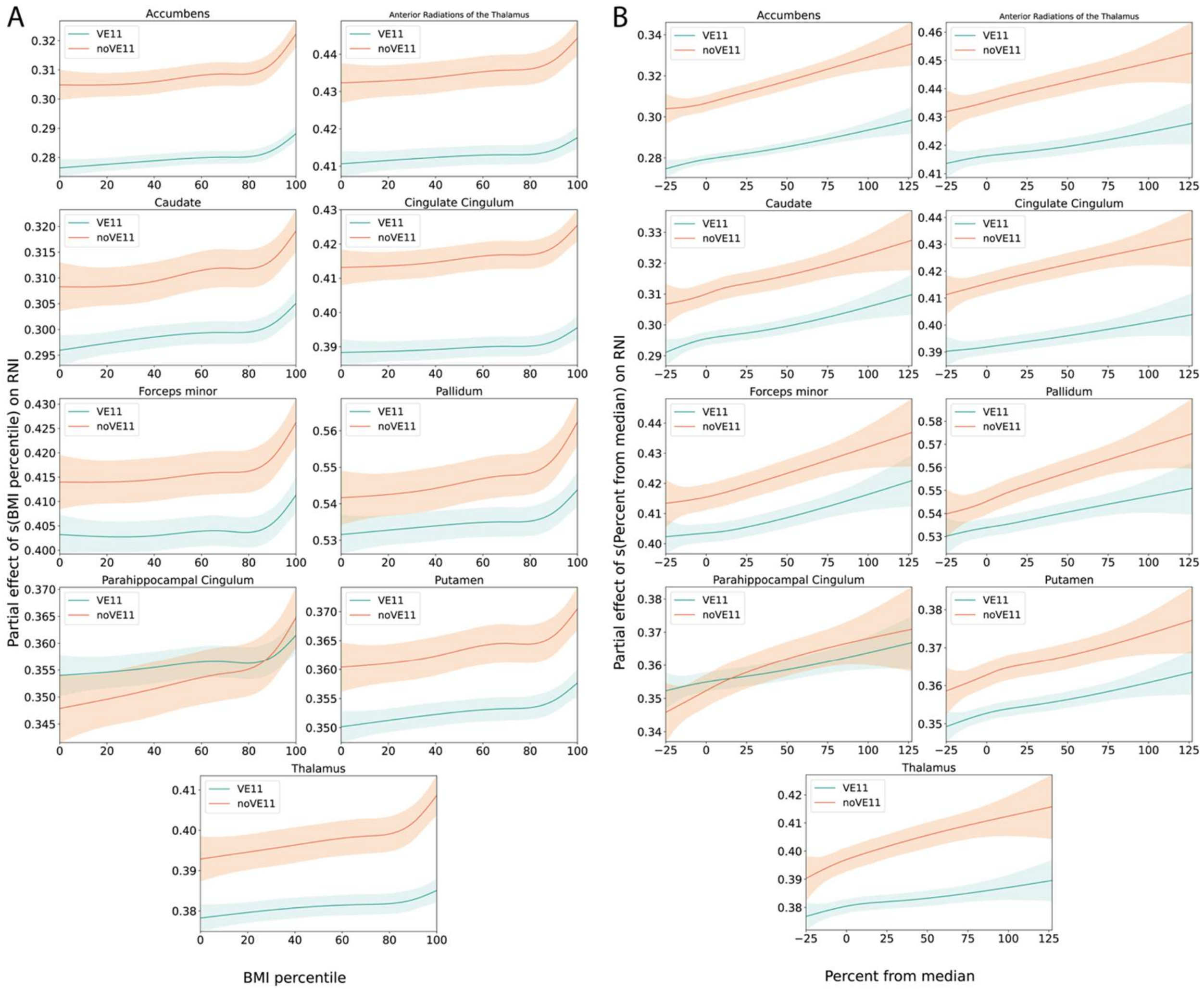


**Note. Scanner-software-group-stratified BMI-RNI associations in selected regions of interest.** Curves show the region-averaged contribution of the BMI spline (plus intercept) on the restricted normalized isotropic signal fraction (RNI) as a function of (A) CDC extended BMI percentile and (B) percent from median BMI, among voxels with an FDR-significant BMI-RNI association. Shaded bands indicate averaged voxelwise 95% confidence intervals. Models were estimated separately in samples with VE11 and non-VE11 scans.

**eFigure 13: Scatterplots of BMI Percentile Beta Coefficient Estimates Between Scanner Software Group**

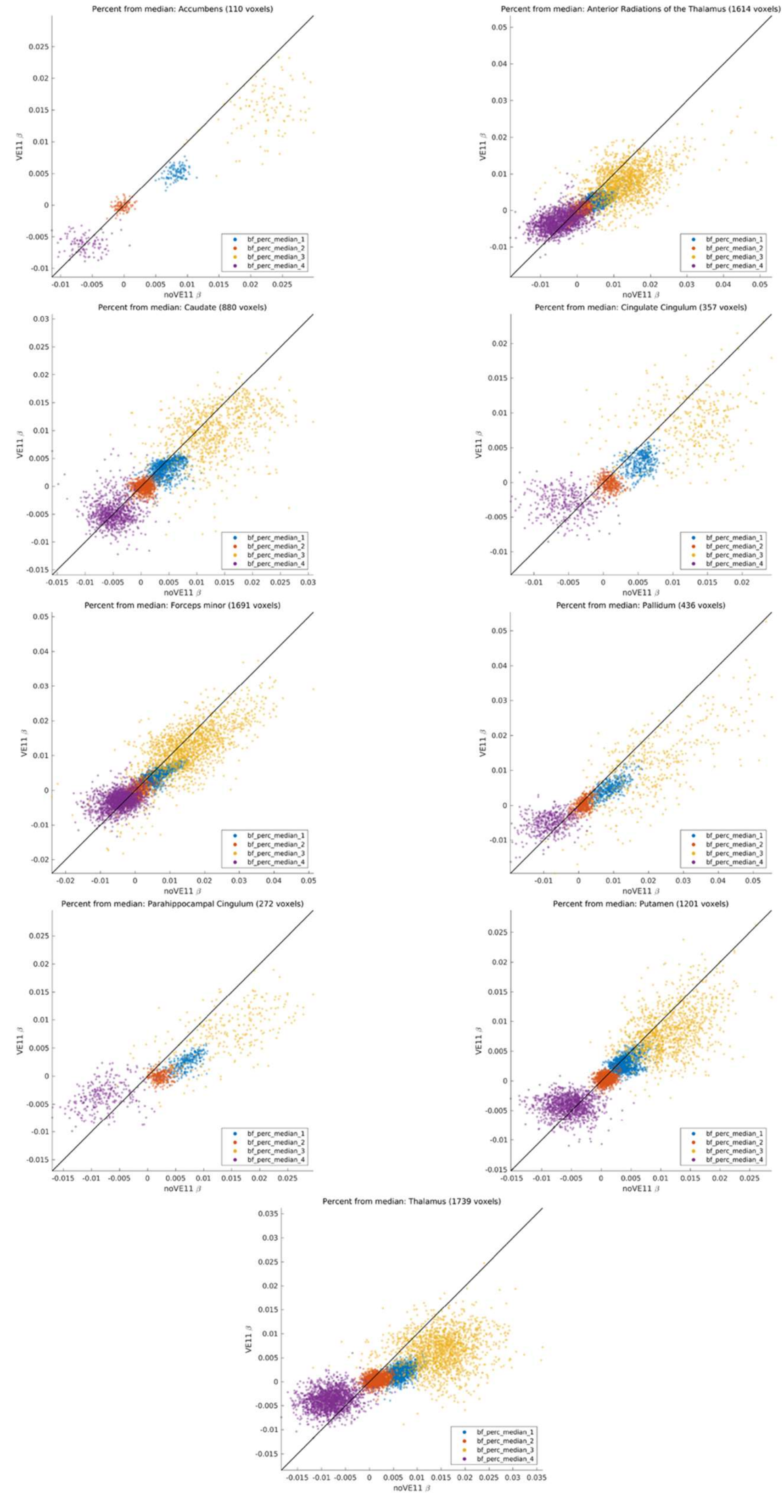

**Note. Scatterplots of BMI Percentile Beta Coefficient Estimates Between Scanner Software Group.** Plots show the correlation between the beta estimates for the basis functions of BMI percentile for VE11 and non-VE11 models from voxels within selected regions of interest.

.

**eFigure 14: Scatterplots of Percent from Median Beta Coefficient Estimates Between Scanner Software Group**

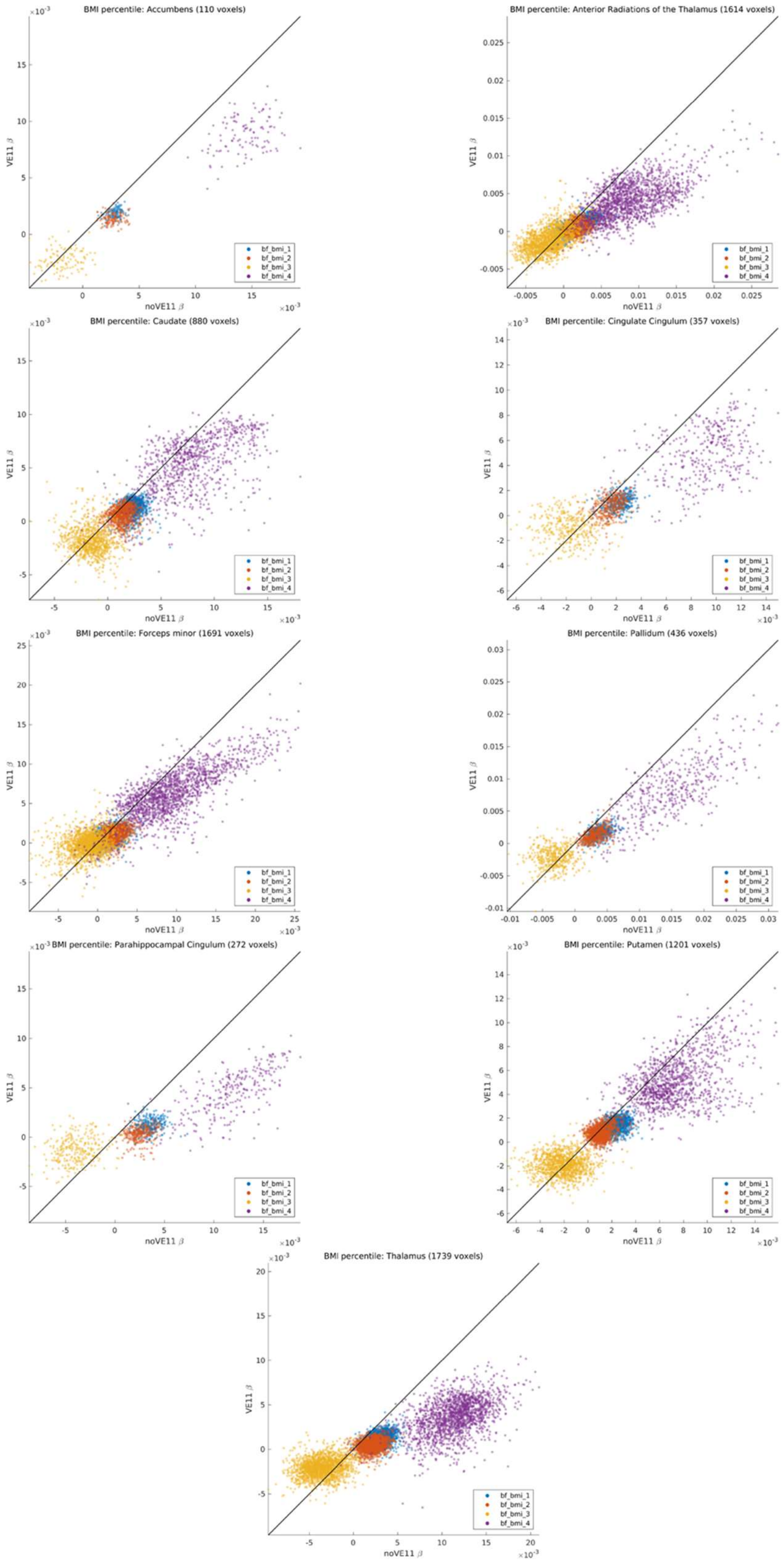

**Note. Scatterplots of Percent from Median BMI Beta Coefficient Estimates Between Scanner Software Group.**

Plots show the correlation between the beta estimates for the basis functions of percent from median BMI for VE11 and non-VE11 models from voxels within selected regions of interest.

**eFigure 15: Scatterplots of Age Beta Coefficient Estimates for BMI Percentile Models Between Scanner Software Group**

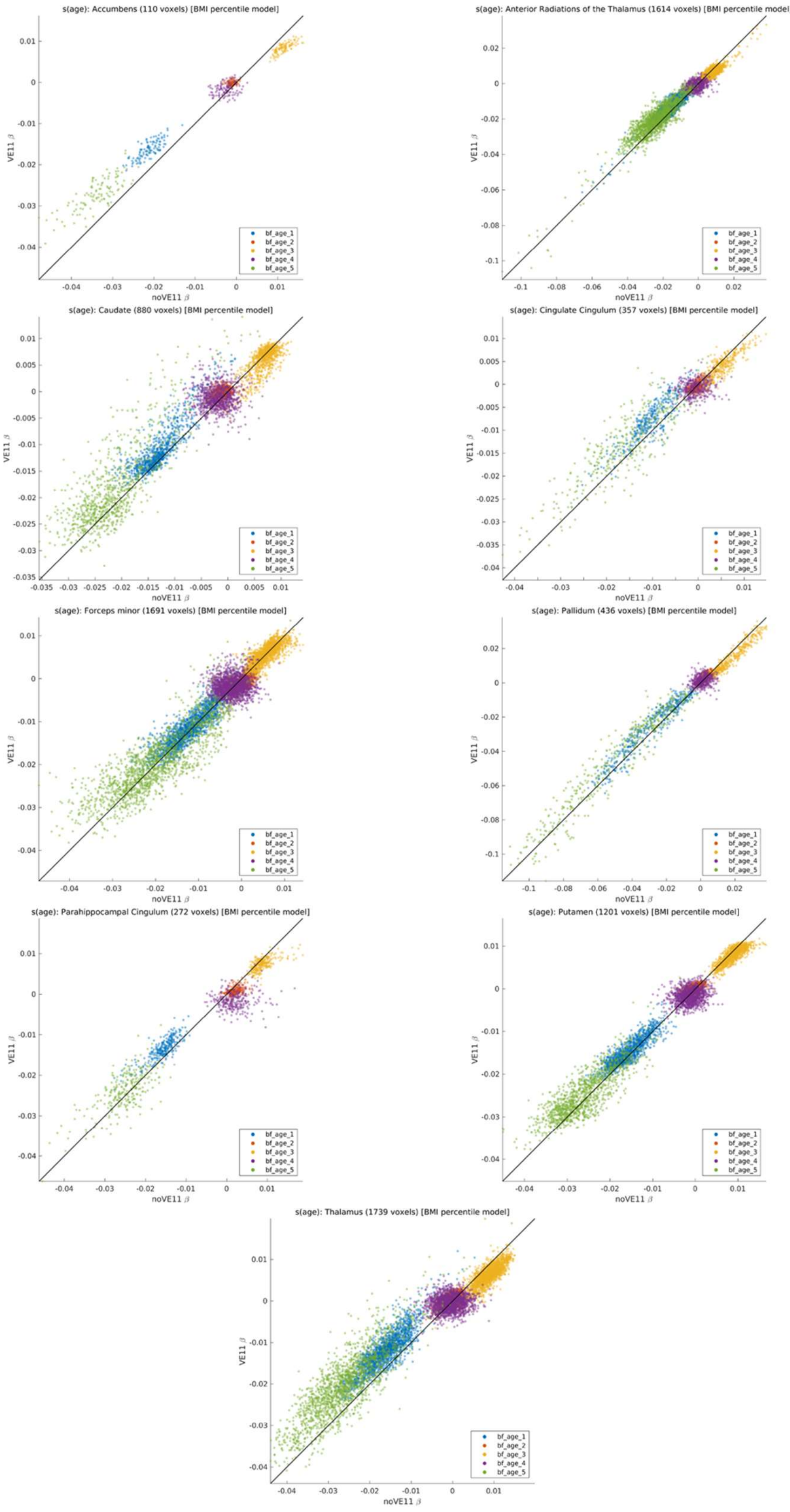

**Note. Scatterplots of Age Beta Coefficient Estimates for BMI Percentile Models Between Scanner Software Group.**

Plots show the correlation between the beta estimates for the basis functions of age for VE11 and non-VE11 models of BMI percentile from voxels within selected regions of interest.

**eFigure 16: Scatterplots of Age Beta Coefficient Estimates for Percent from Median Models Between Scanner Software Group**

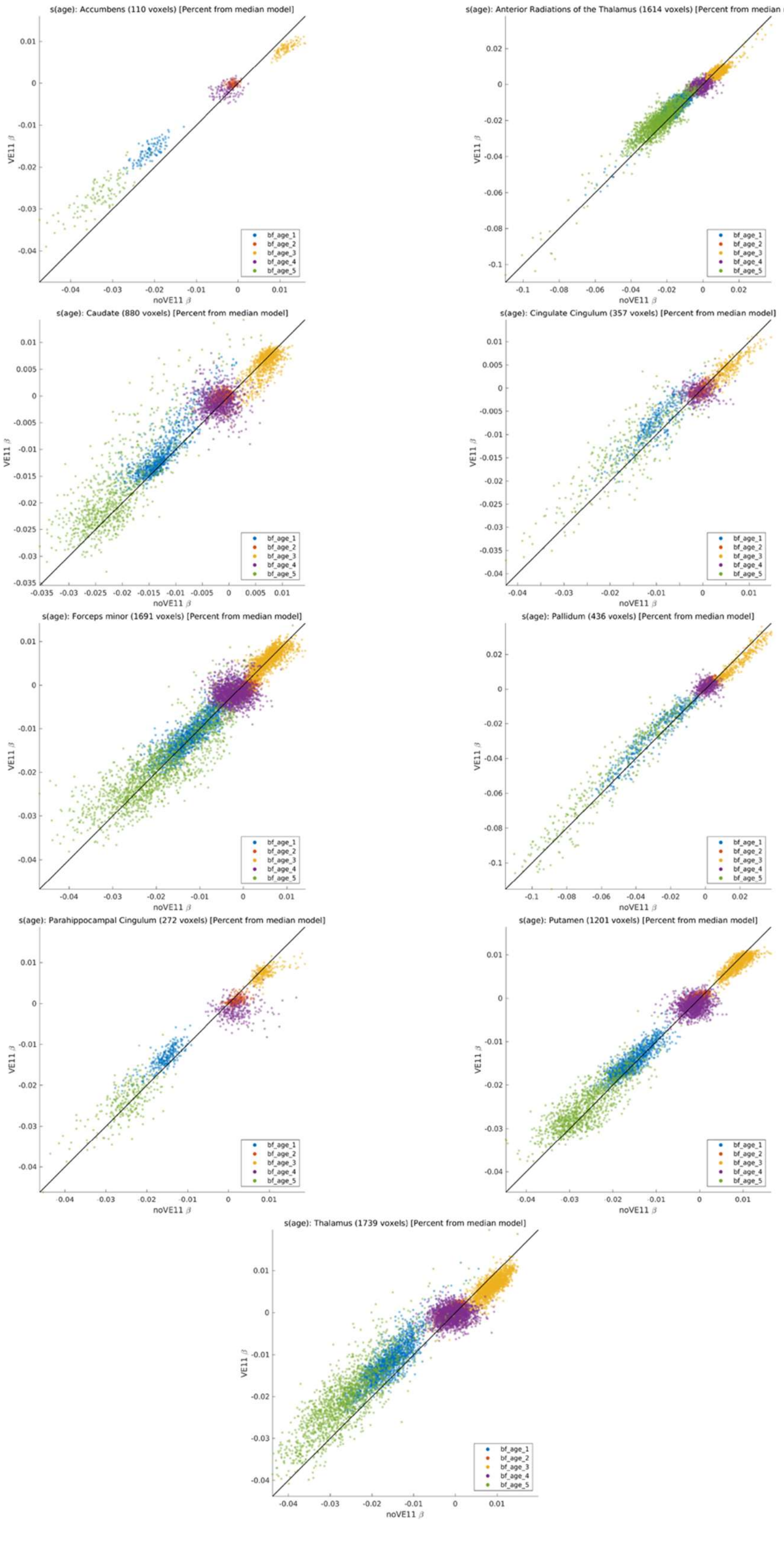

**Note. Scatterplots of Age Beta Coefficient Estimates for Percent from Median BMI Models Between Scanner Software Group.**
Plots show the correlation between the beta estimates for the basis functions of age for VE11 and non-VE11 models of percent from median BMI from voxels within selected regions of interest.
